# Faunistic study on the freshwater ciliates from Delhi, India

**DOI:** 10.1101/2020.07.06.189001

**Authors:** Jeeva Susan Abraham, Renu Gupta, Sripoorna Somasundaram, Ilmas Naqvi, Swati Maurya, Ravi Toteja, Seema Makhija

## Abstract

The ciliated protist communities show richness as well as uniqueness. This is true both for aquatic and soil ciliates. Delhi region lies in the subtropical semi-arid zone wherein the temperatures are highest in May-June and lowest in January. It also receives its monsoonal rainfall during the month of July-August. Thus, the region offers ideal conditions for the growth and proliferation of aquatic living beings. During the past three decades, a series of investigation has been carried out on the freshwater free-living ciliate fauna from fourteen sites at the river Yamuna and different freshwater bodies in Delhi. Samples were brought to the laboratory, ciliates were identified using live-cell observations and silver staining methods. A large number of Spirotrich species and a lower proportion belonging to class Heterotrichea, Litostomaea, Phyllopharyngea, Oligohymenophorea, Prostomatea, and Colpodea were identified. A total of 55 species belonging to 7 classes, 16 orders, 26 families, 40 genera were identified and documented. Ciliate diversity was found to be highest in the water sample from Okhla bird sanctuary (OBS). All ciliate species recorded during the present study have been listed and their general characteristics have been discussed.

## Introduction

Ciliates constitute an important component of freshwater ecosystem and are the most conspicuous group of eukaryotic unicellular organisms (Corliss 1979). Ciliates form an important link in food webs as they act as primordial component of the microbial loop between micro- and macro-invertebrates in the aquatic ecosystem (Aghaindum & Menbohan 2012; Tarbe *et al.* 2011). The ciliates occur in various kinds of aquatic environments, wherein sufficient nutrients, moisture, and appropriate microhabitats are available (Corliss 2002; Mironova *et al.* 2012). Ciliates are known to be useful to man for many purposes such as the determination of water quality and the description of habitat type within the water column (Debastiani *et al.* 2016; Fenchel 1987; Finlay *et al.* 1997; Foissner 1998; Hou *et al.* 2016; Madoni 2000; Shukla & Gupta 2001; Sivasankar *et al.* 2018). Ciliates with their large size, short life cycle, high reproduction rate and lack of cell wall help in detecting the water, soil quality, and the environmental impacts in a short timescale (Abraham *et al.* 2017; Groliére *et al.* 1990; Somasundaram *et al.* 2019; Toteja *et al.* 2017).

The Indian Subcontinent is rich in biodiversity as it has diverse ecosystems. Despite this richness, data concerning freshwater ciliate diversity from India is rather scarce, few reports based on morphological identifications are available (Abraham *et al.* 2019a; Bhatia 1936; Bharti & Kumar 2019; Bindu *et al.* 2018; Chanda *et al.* 2019; Gupta *et al.* 2002; Elangovan & Gauns 2018; Kalavati & Raman 2008; Mahajan & Nair 1971; Makhija *et al.* 2016; Purushothaman *et al.* 2017; Somasundaram *et al.* 2015; Rakshit & Sarkar 2016). And a few novel genera and species of ciliates have been reported from Delhi region (Arora *et al.* 1999; Bharti *et al.* 2019; Gupta *et al.* 2001, 2003, 2006, 2017, 2020; Kamra *et al.* 1994, 2008; Kamra & Kumar 2010; Kaur *et al.* 2019; Kumar *et al.* 2015; Naqvi *et al.* 2006, 2016; Singh & Kamra 2013, 2015). Knowledge of biodiversity of ciliates is becoming increasingly important for modelling and managing ecosystems as their significant role has become better known (Abraham *et al.* 2019b; Basuri *et al.* 2020; Dopheide *et al.* 2008; Li *et al.* 2010).

The present study has been carried out to identify ciliates from different freshwater habitats in Delhi region, India to reveal the hidden ciliate diversity. This report is a compilation of ciliates investigated and identified over a period of past 30 years using morphological approach. This communication reports a brief diagnosis of the Indian population of ciliate species identified from 14 stations in Delhi which include freshwater from River Yamuna at Hindon canal, Okhla barrage, Okhla bird sanctuary and Wazirabad, lakes (Bhalaswa lake, Sanjay lake and Sarpakar lake in Kamla Nehru Ridge), and man-made ponds (Nand Nagri colony, Nehru vihar, Nirankari colony, Raj Ghat, Rithala, Sarai Kale Khan and Wazirabad). A total of 55 species were identified from 7 classes, 16 order, 26 families, 40 genera and documented. Diversity pattern was highest in Okhla Bird Sanctuary (OBS). The majority of the species reported here belong to the class Spirotrichea.

## Materials and methods

### Study area

Delhi is stretched between 21° 36’ 36“ N and 77° 13’ 48“ E and lies in Northern India. It is bordered by states of Haryana on the north, west and south and Uttar Pradesh (UP) to the east. In the present study, sampling was done from fourteen sites including regions from river Yamuna, lakes, and man-made ponds from Delhi. The details of each site are given in Table 1 and Fig. 1.

**Table 1.** Locations and descriptive features of 14 sampling sites.

| Site | Name of the site | Coordinates<br>(Latitude, Longitude) | General features | Temperature<br>(at the time<br>of collection) | pH (at the<br>time of<br>collection) |
| --- | --- | --- | --- | --- | --- |
| 1. | Hindon canal | 28° 58' 53.49" N,<br>77° 29' 13.36" E | Tributary of river<br>Yamuna into<br>Hindon River | 27 °C | 7–8 |
| 2. | Okhla barrage | 28° 32' 29.79" N,<br>77° 18' 59.04" E | Water from river<br>Yamuna | 26 °C | 6–7 |
| 3. | Okhla Bird Sanctuary | 28° 34' 12" N,<br>77° 18' 8.28" E | Bird sanctuary at<br>the Okhla Barrage<br>over river<br>Yamuna | 30 °C | 7–8 |
| 4. | Yamuna water at<br>Wazirabad | 28° 42' 39.96" N,<br>77° 14' 8.16" E | Yamuna river | 26 °C | 7–8 |
| 5. | Bhalaswa lake | 28° 44' 26.52" N,<br>77° 10' 14.16" E | Clean and clear<br>water, natural<br>freshwater lake | 19 °C | 6–7 |
| 6. | Sanjay Lake | 28° 36' 51.12" N,<br>77° 18' 14.04" E | Artificial lake | 30 °C | 7–7.5 |
| 7. | Sarpakaar lake in Kamla<br>Nehru Ridge | 28° 40' 55.56" N,<br>77° 13' 3.72" E | Artificial lake | 26 °C | 6–7 |
| 8. | Nandnagri Colony | 28° 41' 50.64" N,<br>77° 18' 32.04" E | Temporary pond | 27 °C | 6–7 |
| 9. | Nehru Vihar | 28° 42' 37.44" N,<br>77° 13' 6.24" E | Man-made pond | 25 °C | 6–7 |
| 10. | Nirankari colony | 28° 43' 7.86" N,<br>77° 12' 45" E | Temporary pond | 25 °C | 6–7 |
| 11. | Raj Ghat | 28° 38' 26.16" N,<br>77° 14' 57.84" E | Man-made pond | 34 °C | 7–8 |
| 12. | Rithala | 28° 72' 49.44" N,<br>77° 10' 53.74" E | Man-made pond | 31 °C | 7–8 |
| 13. | Sarai Kale Khan | 28° 35' 36.24" N,<br>77° 15' 23.4" E | Temporary pond | 25 °C | 6–7 |
| 14. | Wazirabad | 28° 43' 19.56" N,<br>77° 13' 5.88" E | Man-made Pond | 26 °C | 7–8 |

**FIGURE 1.**
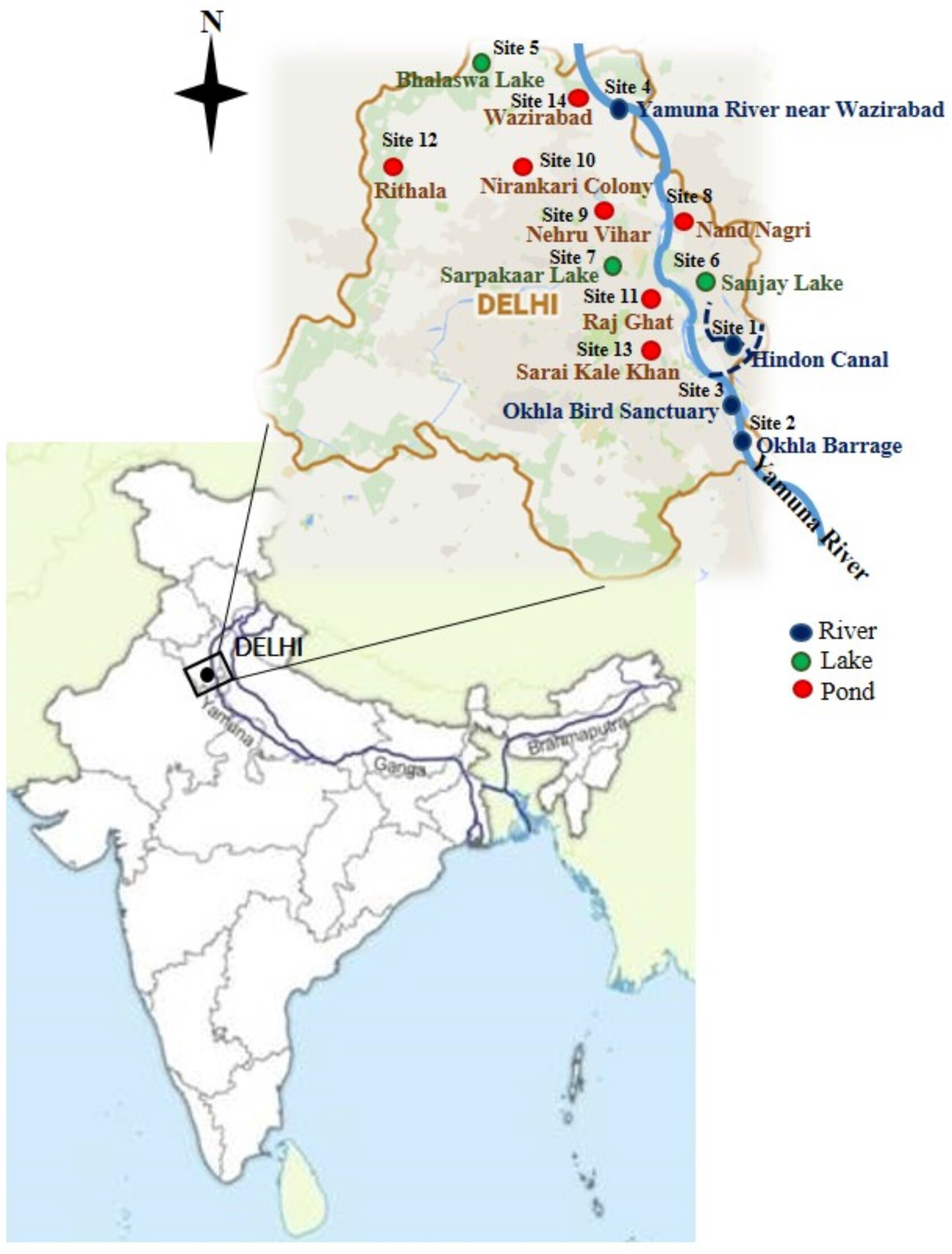
Location and map of sampling sites (site 1–14) in Delhi, India.

### Collection of Samples

Water samples were collected from sites mentioned in table 1. Ecological parameters such as temperature and pH were analyzed on site. Samples were brought to the laboratory, filtered immediately using Nytex nets to remove copepods, crustaceans, debris, and other unwanted materials. Water samples were maintained at room temperature for 3–4 days, with the addition of freshly boiled cabbage pieces to promote the growth of bacteria, which serve as the primary food organism. Samples were subjected to periodic microscopic examination on regular basis.

### *In-vitro* culturing of ciliates

Ciliate from the mixed planktonic cultures were identified under stereoscopic, phase contrast and differential interference contrast microscopes. Single cells were isolated and clonal cultures were raised (wherever possible), maintained and grown in Pringsheim’s medium (Chapman-Andresen 1958) at 22 °C ± 1 °C with boiled cabbage pieces to promote bacterial growth which served as a food source for the ciliates.

### Morphological studies

Live cells were observed under high-power oil immersion objective with bright field, phase contrast and were complemented with protargol staining (Abraham *et al.* 2019b) and Feulgen staining (Feulgen & Rossenbeck 1924; Gupta *et al.* 2018). Measurements were done using eyepiece micrometer or measurement software with proper calibration. Classification, identification, nomenclature, and terminology of ciliate species were done according to Adl *et al.* (2019), Berger (1999), Corliss (1979), Foissner (1998), Lynn & Small (2000), and Lynn (2008).

## Results and discussion

### Ciliate diversity

A total of 55 species belonging to 7 classes, 16 orders, 26 families and 40 genera were identified from freshwater habitats in the Delhi region (Figs 2–6, Table 2 and 3). The taxa belonged to seven classes as follows: Spirotrichea (twenty three taxa), Heterotrichea (four taxa), Litostomatea (five taxa), Oligohymenophorea (five taxa), Phyllopharyngea (one taxon), Prostomatea (one taxon), and Colpodea (one taxon) (Table 3).

**FIGURE 2.**
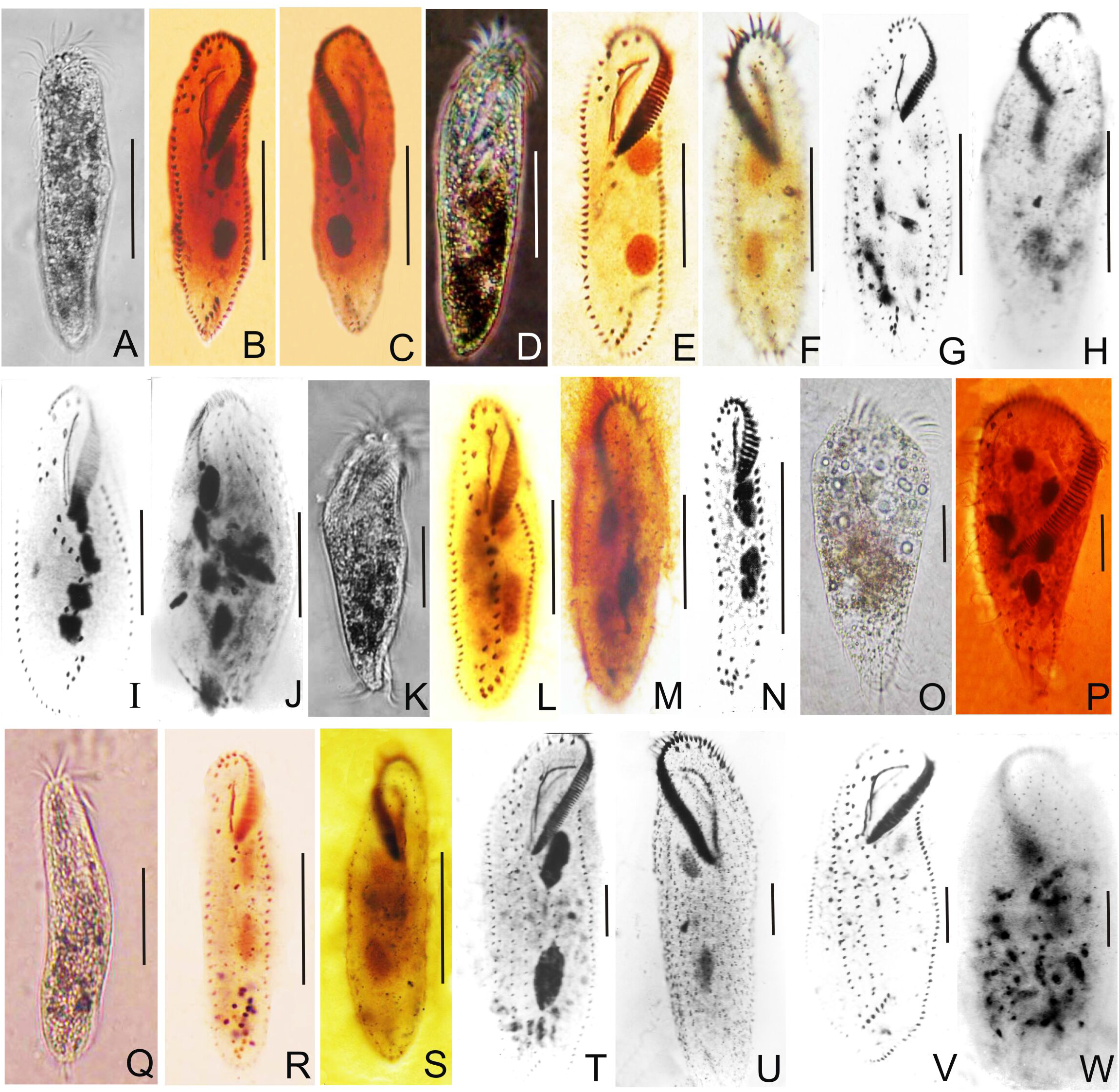
Photomicrographs of freshwater ciliates, **A-C.** *Aponotohymena australis,* **D-F.** *Aponotohymena isoaustralis,* **G-H.** *Architricha indica,* **I-J.** *Gastrostyla steinii,* **K-M.** *Gastrostyla* sp., **N.** *Hemiurosomoida* sp., **O-P.** *Laurentiella* sp., **Q-S.** *Oxytricha granulifera,* **T-U.** *Paraurostyla coronata,* and **V-W.***Paraurostyla weissei.* Scale bars: A-H= 50 μm, I-W= 30 μm.

**FIGURE 3.**
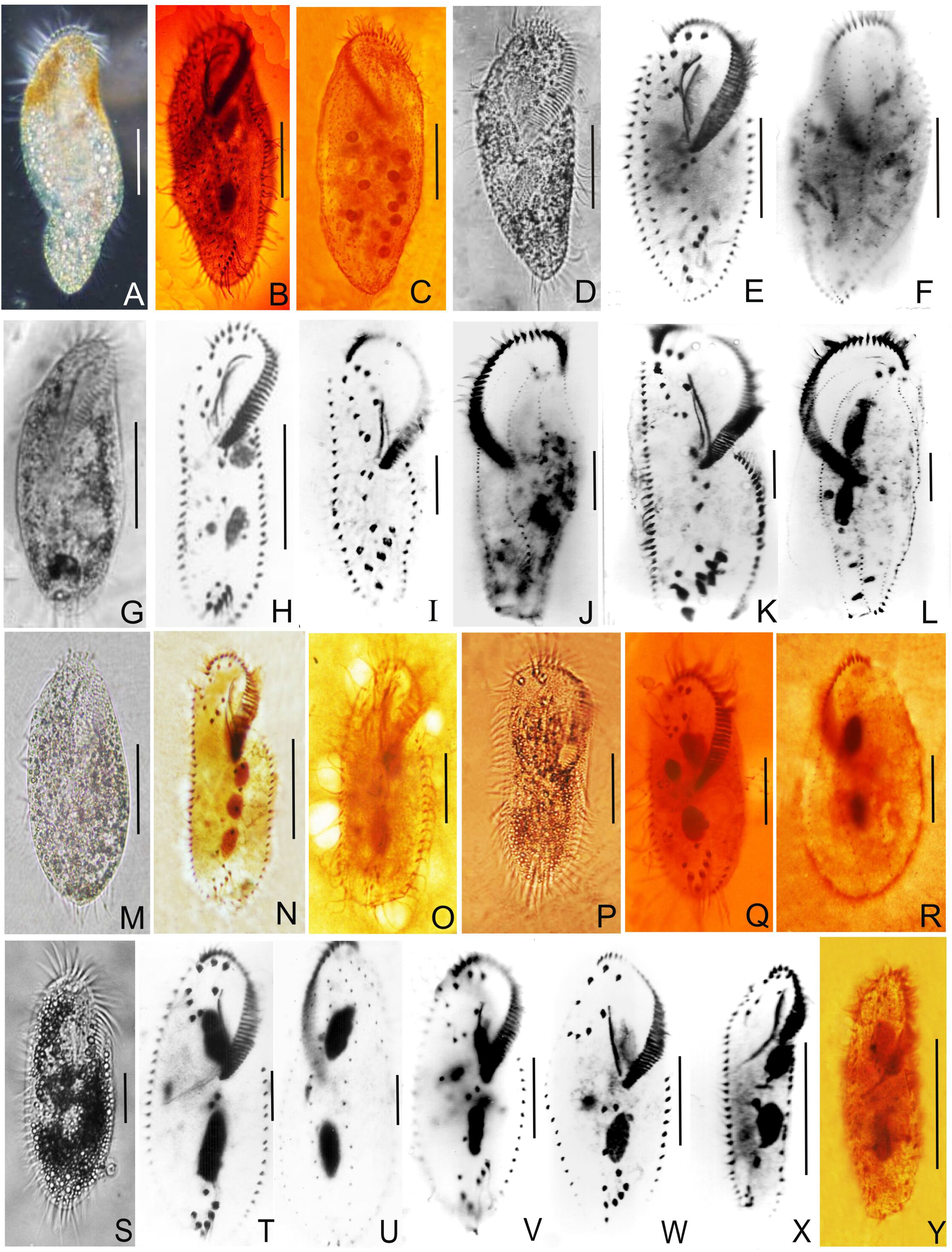
Photomicrographs of freshwater ciliates, **A-C.** *Paraurostyla* sp., **D-F.** *Pleurotricha curdsi,* **G-H.** *Rubrioxyticha indica,* **I-J.** *Stylonychia ammermanni,* **K-L.** *Stylonychia lemnae,* **M-N.** *Sterkiella tetracirrata,* **O.** *Sterkiella* sp., **P-R.** *Tetmemena pustulata,* **S-U.** *Tetmemena saprai,* **V.** *Tetmemena vorax,* **W.** *Tetmemena* sp., **X.** *Urosomoida* sp., **Y.** *Gonostomum* sp. Scale bars: A-F= 50 μm, G-Y= 30 μm.

**FIGURE 4.**
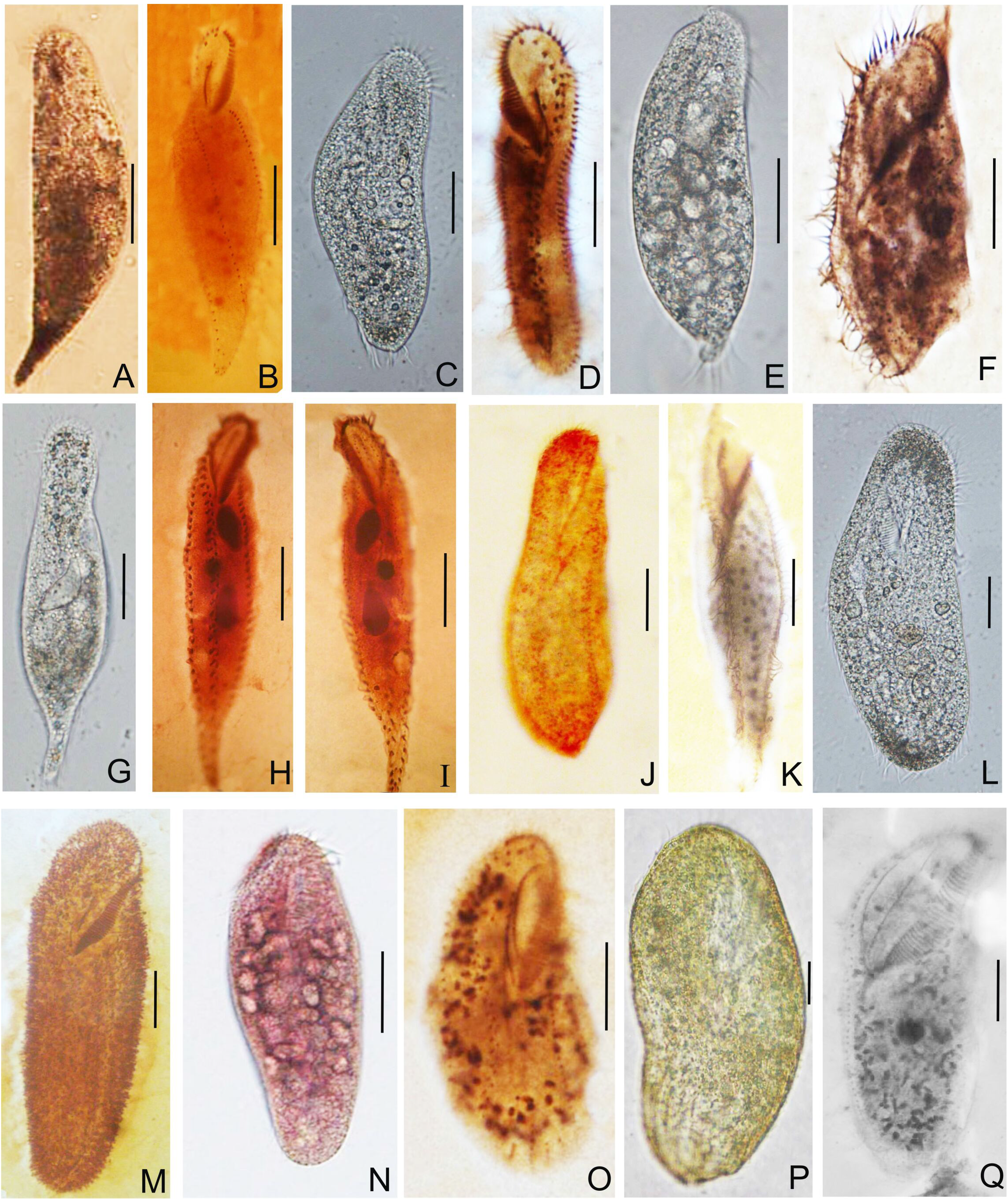
Photomicrographs of freshwater ciliates, **A-B.** *Hemiamphisiella terricola,* **C-D.** *Anteholosticha monilata,* **E-F.** *Uroleptus gallina,* **G-I.** *Uroleptus longicaudatus,* **J-K.** *Pseudokeronopsis* sp., **L-M.** *Pseudourostyla cristata,* **N-O.** *Diaxonella trimarginata,* **P-Q.** *Urostyla grandis.* Scale bars= 30 μm.

**FIGURE 5.**
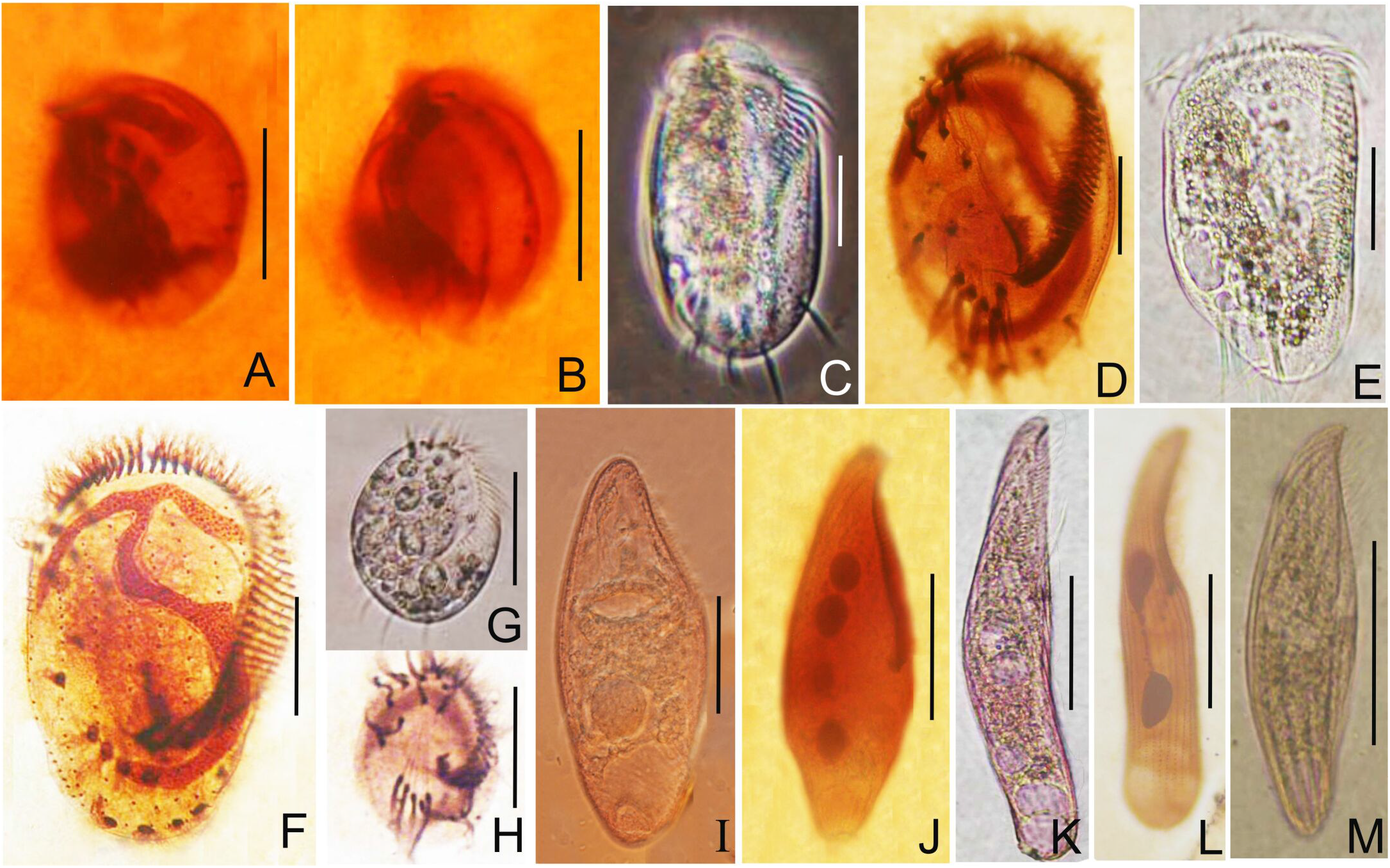
Photomicrographs of freshwater ciliates, **A-B.** *Aspidisca* sp., **C-D.** *Euplotes aediculatus,* **E-F.***Euplotes woodruffi,* **G-H.** *Euplotes* sp. **I-J.** *Blepharisma sinosum,* **K-L.** *Blepharisma undulans,* **M.** *Blepharisma* sp. Scale bars: A, B= 10 μm, C-M= 30 μm.

**FIGURE 6.**
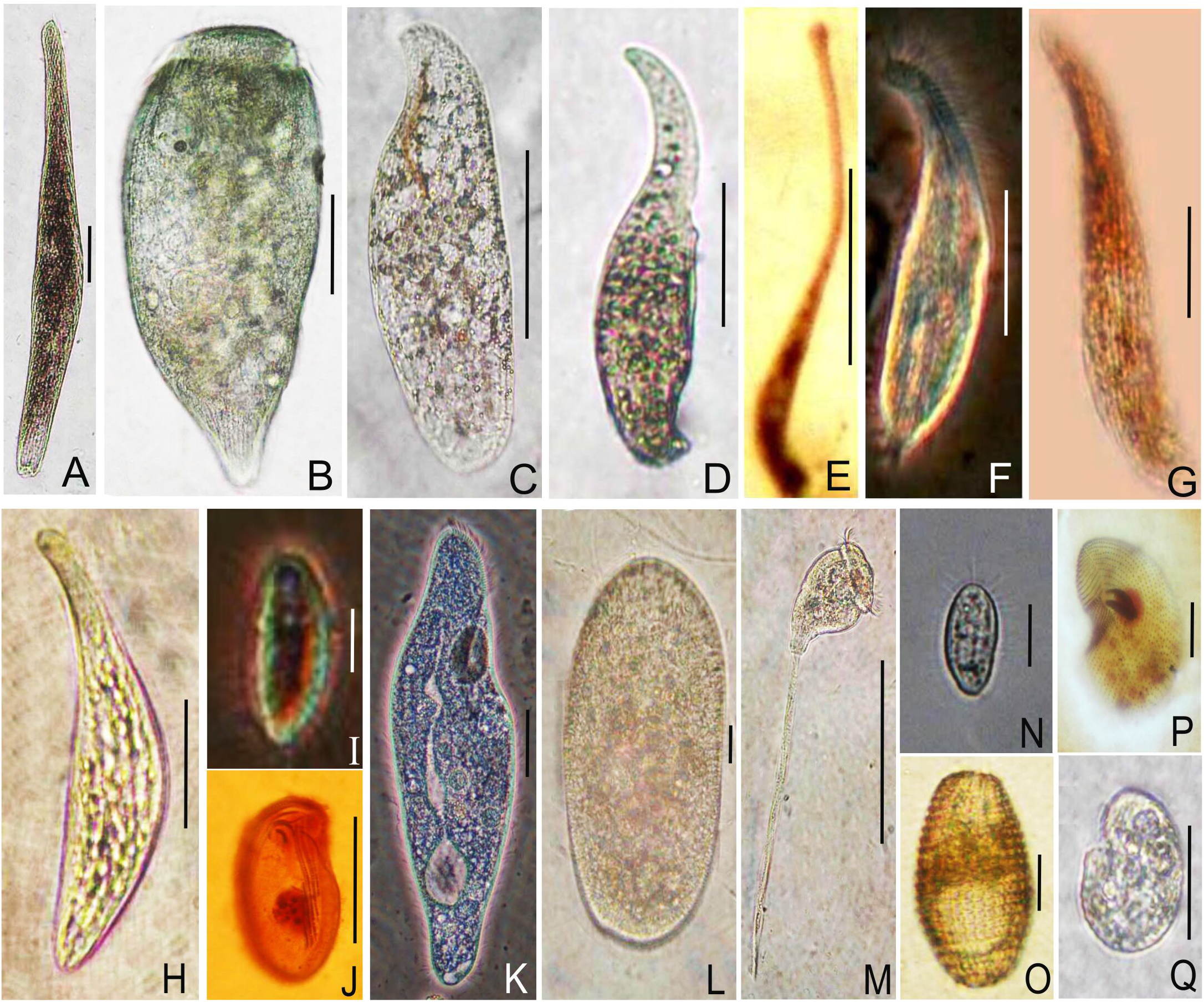
Photomicrographs of freshwater ciliates, **A.** *Spirostomum minus,* **B.** *Stentor* sp. **C.** *Loxodes* sp., **D.** *Dileptus* sp., **E.** *Lacrymaria* sp., **F.** *Loxophyllum* sp., **G.** *Litonotus* sp., **H.** *Spathidium* sp. **I.** *Chilodonella* sp. **J.** *Cyclidium* sp. **K.** *Paramecium multimicronucleatum,* **L.** *Frontonia* sp., **M.** *Vorticella* sp., **N.** *Dexiotricha* sp., **O.** *Coleps* sp., **P.** *Colpoda magna* **Q.** *Colpoda* sp. Scale bars: A-H= 50 μm, I= 10 μm, J-Q= 30 μm.

**Table 2.**
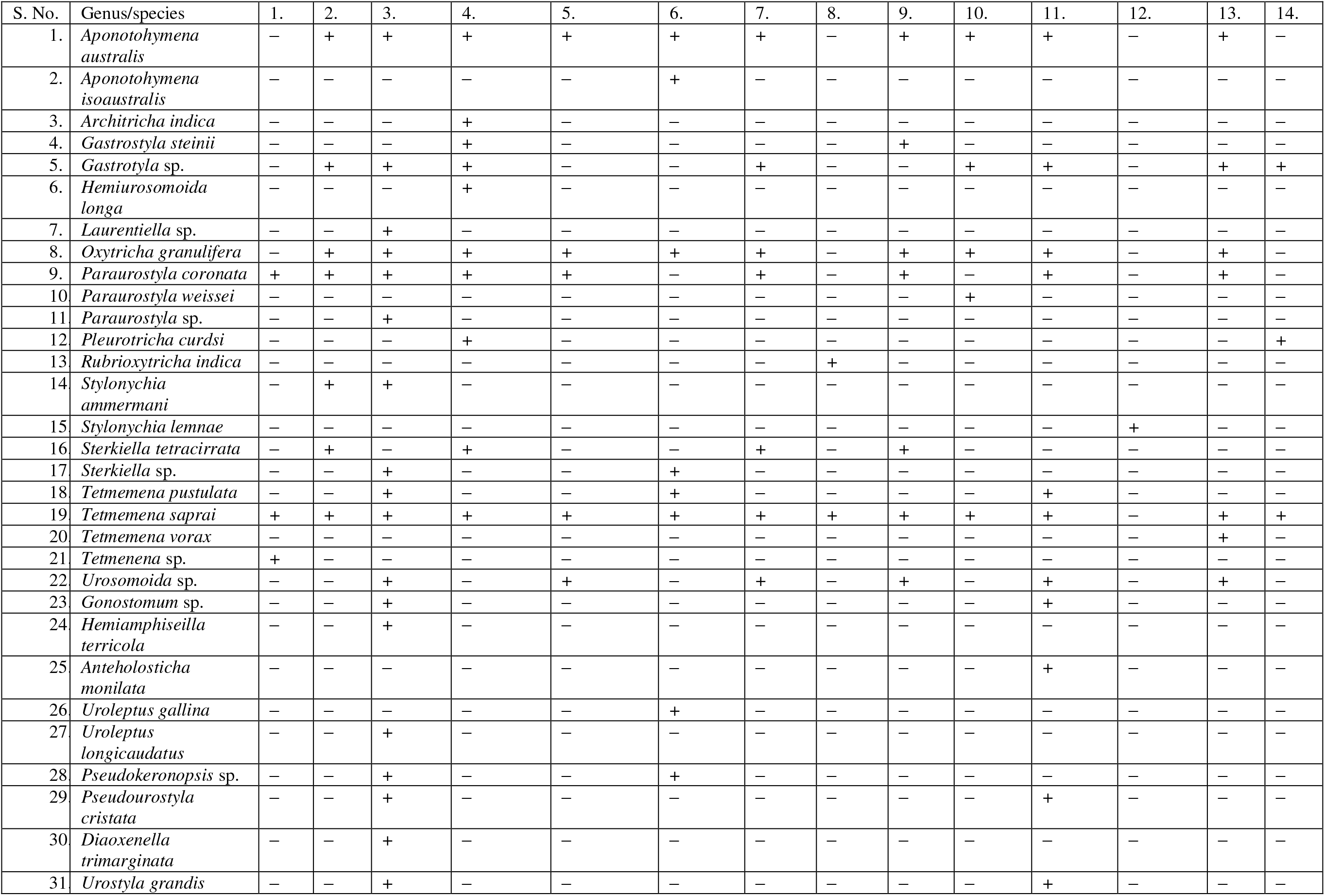
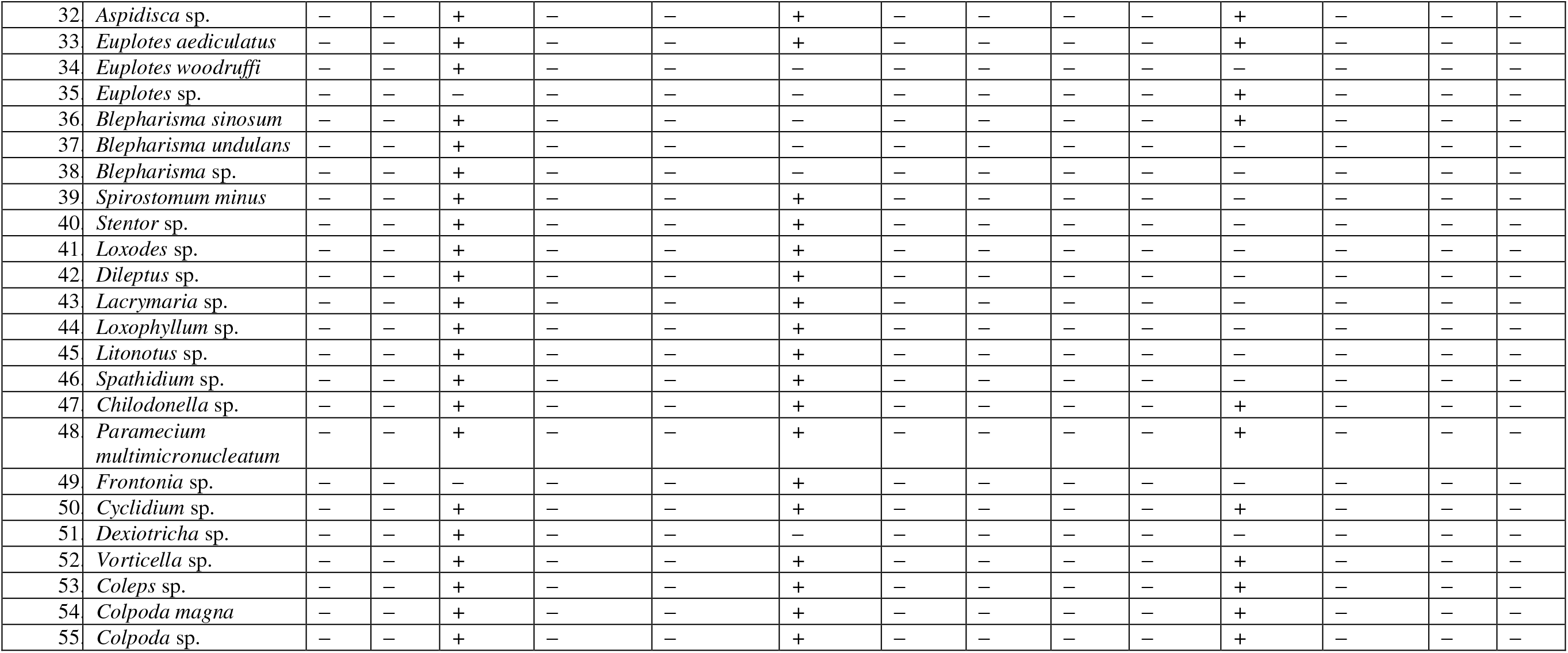
Distribution of 55 free living ciliate species in 14 freshwater bodies in Delhi.

**Table 3.**
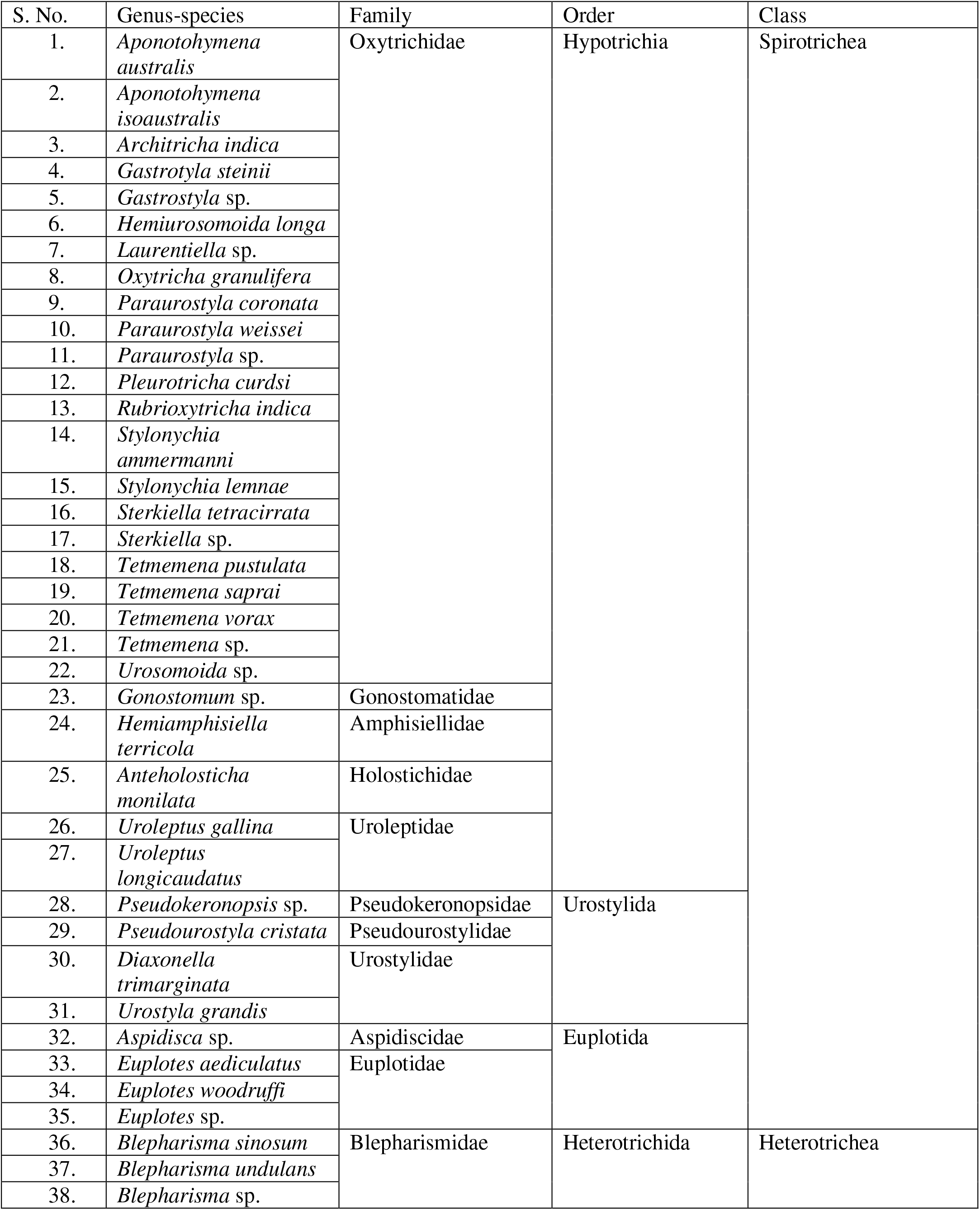
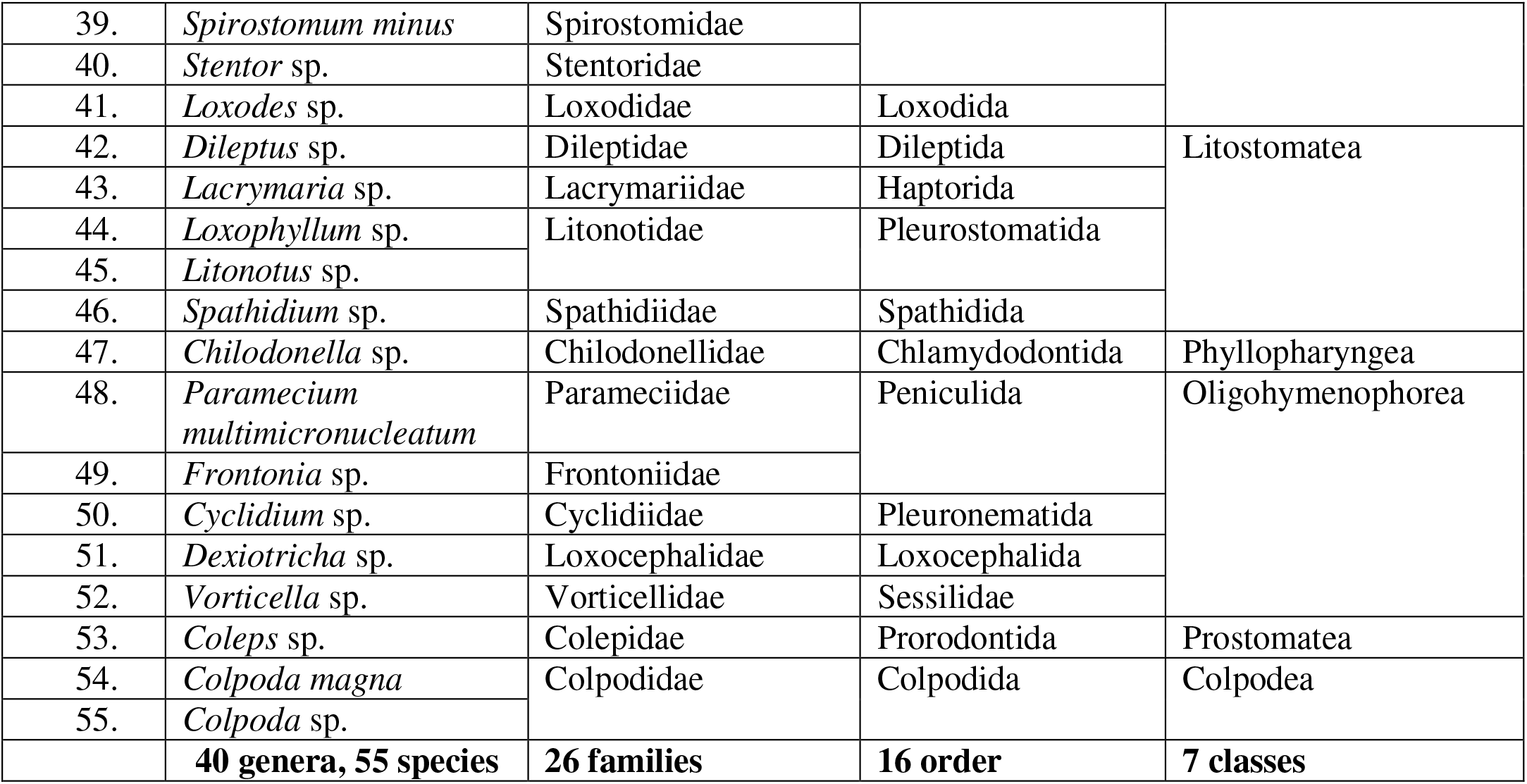
List of 55 free living ciliate species isolated from 14 freshwater habitats in Delhi region (Classification according to Adl *et al.,* 2019).

Class Spirotichea Bütschli, 1889
Order Hypotrichia Stein, 1859
Family Oxytrichidae Ehrenberg, 1830
Genus *Aponotohymena* Foissner, 2016

#### 1. *Aponotohymena australis* Foissner & O’ Donoghue, 1989

Size in vivo 120–150 × 30–40 μm, after protargol staining 128 × 31 μm, length to width ratio 4:1; body flexible, colourless, prolate ellipsoidal, dorsoventrally flattened, subpellicular granules located mainly along the rows of cirri and dorsal kineties, arranged in small groups imparting yellowish orange colouration to the cell; AZM occupies about 33% of the body length with about 44 membranelles, buccal cavity large and deep with anterior end curved; undulating membrane (UMs) with a characteristic hook at the distal end (*Notohymena* pattern); two macronuclei, six to eight micronuclei; 18 frontal-ventral-transverse (FVT) cirri which include three frontal, one buccal, four frontoventral, three postoral ventral, two pretransverse, five transverse cirri (arranged in a linear row); one left and one right row of marginal cirri non confluent posteriorly with an average of 40 and 38 cirri, respectively; six rows of dorsal kineties (DK_1-4_ and DM_1 & 2_), DK_1_–DK_3_ extend along the entire body length, DK_4_ shortened posteriorly, DM_1_ & DM_2_ present in the anterior half of the body, seven to ten caudal cirri at the posterior ends of DK_1, 2 & 4_, DK_1 & 2_ bear two to three caudal cirri each while DK_4_ has three to four caudal cirri at the posterior end; encystment frequent, conjugation observed.

#### 2. *Aponotohymena isoaustralis* Gupta *et al.,* 2017

Size in vivo 140–150 × 40–50 μm, after protargol impregnation 132 × 34 μm, body length to width ratio about 4:1; body elongated, dorsoventrally flattened, flexible, cytoplasm colourless with greenish appearance at the margins; adoral zone of membranelles (AZM) occupies about 34% of the body length with an average of 36 membranelles; undulating membrane (UM) with a characteristic hook at the distal end (*Notohymena* pattern); two to four ellipsoidal macronuclei, amicronucleate; 18 FVT cirri with three frontal, one buccal, four frontoventral, three postoral ventral, two pretransverse ventral, five transverse cirri; one left and one right row of marginal cirri non confluent posteriorly with an average of 34 and 32 cirri, respectively; six rows of dorsal kineties (DK_1-4_ and DM_1-2_), seven caudal cirri (constant) in 2+2+3 pattern, at the posterior ends of kineties DK_1,2 & 4_; generation time under laboratory conditions 11 h, encystment frequent, conjugation not observed.

Genus *Architricha* Gupta *et al.,* 2006

#### 3. *Architricha indica* Gupta *et al.,* 2006

Size in vivo about 130–140 × 40–50 μm, after protargol impregnation 122 × 43 μm, length to width ratio 3:1, oval shaped, dorsoventrally flattened, flexible, cytoplasm colorless, yellow– green cortical granules in short linear rows; AZM occupies about 30% of the body length with an average of 34 membranelles; UMs in *Oxytricha* pattern; two ellipsoidal macronuclei, two spherical micronuclei; 18 FVT cirri which includes three frontal, one buccal, four frontoventral, three postoral ventral, two pretransverse, five transverse cirri (arranged in tick mark pattern); two left and three right rows of marginal cirri; six rows of dorsal kineties (DK_1–4_ and DM_1–2_), DK_4_ short and curved; three caudal cirri, one each at the posterior ends of kineties DK_1, 2 & 4_; cells encystement and excystment frequent, conjugation not observed.

Genus *Gastrostyla* Engelmann, 1862

#### 4. *Gastrostyla steinii* Engelmann, 1862

Size in vivo 90–100 × 40–50 μm, after protargol staining 90 × 45 μm, length to width ratio 2:1; flexible, dorsoventrally flattened, oval, anterior portion narrow, posterior portion broad; AZM occupies about 40% of the body length with an average of 45 membranelles; four ovoid macronuclei, two to four spherical micronuclei; 23–26 FVT cirri; ventral cirri variable in numbers ranging between 14–17, arranged as a median cirral row, one left and one right row of marginal cirri with an average of 24 and 28 cirri, respectively; six rows of dorsal kineties (DK_1-4_ and DM_1-2_), DK_3_ and DK_4_ shortened anteriorly; three caudal cirri, one each at the posterior ends of kineties DK_1, 2 & 4_, second caudal cirrus usually placed slightly above the level of the other two caudal cirri; generation time under laboratory conditions 9 ± 0.5 h, encystment, excystment, cannibalism and reorganization frequent, conjugation not observed.

#### 5. *Gastrostyla* sp

Size in vivo about 95–100 × 30–32 μm, after protargol staining 74–100 × 21–35 μm, length to width ratio 3:1; flexible, dorsoventrally flattened; AZM occupies 37% of the body length with an average of 32 membranelles; undulating membranes in *Oxytricha* pattern; two ovoid macronuclei, two to six spherical micronuclei; 24–30 frontoventral transverse cirri, number of frontoventral and postoral ventral cirri is distinctly more than seven and they form a continuous slightly oblique frontoventral row, two pretransverse, five transverse cirri, one left and one right marginal row non confluent with an average of 27 and 30 cirri; six rows of dorsal kineties (DK_1-4_ and DM_1-2_), DK_3_ shortened posteriorly, DK_4_ commences subequatorially, three to four caudal cirri at the posterior ends of kineties DK_1, 2 & 4;_ encystment and excystment frequent, conjugation not observed.

Genus *Hemiurosomoida* Gelei & Szabados, 1950

#### 6. *Hemiurosomoida longa* Gelei & Szabados, 1950

Size in vivo 50–60 × 15–20 μm, after protargol impregnation about 50 × 12 μm, length to width ratio 4:1; flexible, dorsoventrally flattened, cells extremely thin with narrowly rounded ends; AZM occupies 30% of the body length with an average of 18 membranelles; undulating membranes in *Oxytricha* pattern; two macronuclei, two to ten spherical micronuclei; 17 FVT cirri with three frontal, one buccal, four frontoventral, three postoral ventral, two pretransverse, four transverse cirri, one left and one right row of marginal cirri with 16 cirri in each row on an average; four rows of dorsal kineties (DK_1-3_ and DM_1_), two caudal cirri; encyst frequent, conjugation not observed.

Genus *Laurentiella* Dragesco & Njine, 1971

#### 7. *Laurentiella* sp

**S**ize in vivo 150–180 × 80–100 μm, after protargol impregnation about 151 × 74 μm, length to width ratio 2:1; body stiff, almost triangular, oral cavity wide and expansive, anterior end broad and posterior end tapered; AZM occupies about 50% of body length with about an average of 48 membranelles, distal end distinctly overlapping on the right body margin; undulating membranes (UMs) in typical *Oxytricha* pattern; four to six macronuclei, seven to eight micronuclei; four cirral rows including one buccal cirrus, about nine to ten cirri arranged in cirral rows 2–3, five transverse cirri; one left and one right row of marginal cirri with an average of 24 and 36 cirri, respectively; six rows of dorsal kineties (DK_1–4_ and DM_1 & 2_), three caudal cirri; encystment frequent, conjugation not observed.

Genus *Oxytricha* Ehrenberg, 1838

#### 8. *Oxytricha granulifera* Foissner & Adam, 1983

Size in vivo 68–109 × 22–25 μm, after protargol impregnation 71 × 16 μm on an average, length to width ratio 4.4:1; body flexible, lanceolate anterior and posterior ends, entire cell body loaded with non-pigmented granules which stain heavily with protargol; AZM occupies about 30% of the body length with an average of 25 membranelles; UMs in typical *Oxytricha* pattern; two macronuclei, two micronuclei; 18 FVT cirri which includes four frontal, one buccal, four frontoventral, three postoral ventral, two pretransverse, five transverse cirri arranged in an oblique linear row; one left and one right row of marginal cirri with an average of 22 and 23 cirri, respectively, non confluent posteriorly; six rows of dorsal kineties (DK_1–4_ and DM_1–2_); encystement frequent, conjugation observed.

Genus *Paraurostyla* Borror, 1972

#### 9. *Paraurostyla coronata* Arora *et al.,* 1999

Size in vivo 200–220 × 60–70 μm, size after protargol impregnation 200 × 62 μm, length to width ratio 3:1; body flexible, diffused green coloured appearance, pink coloured anterior and posterior extremities, dorsoventrally flattened; AZM occupies 33% of the cell length with an average of 68 membranelles; UM in typical *Oxytricha* pattern; two macronuclei, four micronuclei; seven frontal cirri, one buccal cirrus, six to eight (usually seven) longitudinal rows of ventral cirri lie on the right half of the ventral surface, V_1_ row short consisting of two to five cirri of which the posterior most cirrus is present in the postoral region, highly variable number of cirri, six to ten transverse cirri form an oblique row; one left and one right row of marginal cirri with an average of 55 and 48 cirri, respectively, non confluent posteriorly; six rows of dorsal kineties (DK_1–4_ and DM_1–2_), DK_1–3_ run through the entire length of the cell, DK_4_ begins at the equatorial region, DM_1 & 2_ short, terminating at different levels above the equatorial region; three rows of caudal cirri containing 15–18 cirri present at the posterior ends of DK_1, 2 &4_ with three to seven cirri in each row; generation time under laboratory conditions 12 ± 0.5 h, frequent cortical reorganization, encystment rare whereas cannibalism quite common, conjugation not observed.

#### 10. *Paraurostyla weissei* Stein, 1859

Size in vivo about 200–220 × 80–100 μm, after protargol staining about 170 × 70 μm, length to width ratio of 2.8:1; body flexible, slender, dorsoventrally flattened, underlying yellow-greenish granules, elliptical, narrowing and sometimes tapering towards the posterior end; AZM occupies 36% of body length with about 60 membranelles; UMs in typical *Oxytricha* pattern; two macronuclei and four micronuclei; six large frontal cirri arranged in two diagonal rows of four and two cirri, five to six longitudinal rows of ventral cirri, number of ventral cirri in each row variable; seven to eight transverse cirri; one left and one right row of marginal cirri with an average of 46 and 48 cirri, respectively; six to eight rows of dorsal kineties, DK_1–3_ complete and cover the entire length, short rows of two to four dorsomarginal rows (DMs); caudal cirri at the posterior ends of DK_1, 2 & 4_, DK_2_ possess fewer caudal cirri while DK_4_ bears more of them; generation time under laboratory conditions 11 ± 0.5 h, reorganization and cannibalism frequent, encystment and conjugation not observed.

#### 11. *Paraurostyla* sp

Size in vivo 200–242 × 52–76 μm, **s**ize after protargol impregnation 171 × 63 μm on an average, length to width ratio 2:1; body flexible, dorsoventrally flattened, diffused green coloured general appearance, yellowish-green coloured anterior and posterior extremities; AZM occupies 37% of the cell length with an average of 55 membranelles; UM in typical *Oxytricha* pattern; two macronuclei, six to eight micronuclei; six frontal cirri, one buccal cirrus, five to eight (usually six) longitudinal rows of ventral cirri lie on the right half of the ventral surface, short row of V_1_ containing two to four cirri, highly variable number of cirri in each of ventral row ranging from 12–22 cirri on an average in each row, seven to eight transverse cirri form an oblique row; one left and one right row of marginal cirri with an average of 44 cirri in each row, non confluent posteriorly; six rows of dorsal kineties (DK_1–4_ and DM_1–2_), DK_1-3_ run through the entire length of the cell, DK_4_ begins at the equatorial region, DM_1 & 2_ short, terminating above the equatorial region; three rows of caudal cirri containing seven to 12 cirri present at the posterior ends of DK_1, 2 & 4_ with three to seven cirri in each row; generation time under laboratory conditions 12 ± 0.5 h, frequent cortical reorganization, encystment rare whereas cannibalism quite common, conjugation not observed.

Genus *Pleurotricha* Stein, 1859

#### 12. *Pleurotricha curdsi* Shi *et al.* 2002

Size in vivo about 150–160 × 60–70 μm, after protargol staining about 140 × 64 μm, length to width ratio of 2:1; body rigid, oval shape tapering gradually posteriorly, right margin distinctly convex; AZM covering 40% of the body length consisting of about 50 membranelles on an average; UMs in typical *Oxytricha* pattern; two macronuclei, two micronuclei; 20 FVT cirri with nine frontal cirri, six postoral ventral cirri, five transverse cirri arranged in two groups of three and two cirri; one row of left marginal cirri with an average of 22 cirri and two rows of right marginal cirri with an average of 28 (in RMC1) and 13 cirri (in RMC2); six rows of dorsal kineties (DK_1–4_ and DM_1–2_), DK_1–4_ traverse the entire cell length, DM_1 & 2_ run about half and one fourth of the cell length respectively, three caudal cirri one each at the posterior ends of DK_1, 2 & 4_ placed in continuation with the terminal cirrus of the RMC1; generation time under laboratory conditions 5 ± 0.5 hr, encystment frequent, conjugation not observed.

Genus *Rubrioxytricha* Berger, 1999

#### 13. *Rubrioxytricha indica* Naqvi *et al.,* 2006

Size in vivo 70–80 × 25–30 μm, after protargol impregnation 69 × 27 μm on an average, length to width ratio is 2.6:1; body highly flexible, elliptical in shape, cytoplasm with dark brown crystalline inclusions, spherical dark green cortical granules present in single or clusters of two to five; AZM covering 40% of body length with an average of 29 membranelles; UM in *Cyrtohymena* pattern; two macronuclei, two or three micronuclei; 18 FVT cirri with three frontal, one buccal, four frontoventral, three postoral ventral, two pretransverse, five transverse cirri; one left and one right row of marginal cirri with an average of 19 and 22 cirri, respectively; five rows of dorsal kineties (DK_1–3_ and DM_1–2_), one caudal cirrus; encystment and excystment frequent; conjugation not observed.

Genus *Stylonychia* Ehrenberg, 1830

#### 14. *Stylonychia ammermanni* Gupta *et al.,* 2001

Size in vivo 140–145 × 50–60 μm, after protargol staining is about 135 × 50 μm, length to width ratio of 2.5:1; rigid, broad anterior and narrow posterior; AZM occupies 53 % of the body length with an average of 51 membranelles; UMs in *Stylonychia* pattern; two macronuclei, two to four micronuclei; 18 FVT cirri with three frontal, one buccal, four frontoventral, three postoral ventral, two pretransverse, five transverse cirri, one left and one right row of marginal cirri with an average of 17 and 24 cirri, respectively, non confluent posteriorly; six rows of dorsal kineties (DK_1-4_ and DM_1, 2_), DK_1-3_ curved apically, three caudal cirri one each at the end of dorsal kineties DK_1, 2 & 4_ equally spaced; generation time under laboratory conditions 11 ± 0.5 hrs, encystment rare, conjugation not observed.

#### 15. *Stylonychia lemnae* Ammermann & Schlegel, 1983

Size in vivo 180–190 × 75–85 μm, after protargol impregnation about 188 × 75 μm, length to width ratio of 2.5:1; body rigid, dorsoventrally flattened, anterior and posterior ends are rounded with more or less parallel margins; AZM occupies about 50% of the body length with 62 membranelles on an average; UMs in *Stylonychia* pattern; two macronuclei, two to four micronuclei; 18 FVT cirri with three frontal, one buccal, four frontoventral, three postoral ventral, two pretransverse, five transverse cirri arranged in two groups of three and two; one left and one right row of marginal cirri with 26 and 38 cirri on an average, respectively; six rows of dorsal kineties (DK_1–4_ and DM_1, 2_), DK_1–3_ complete rows and bent apically, the fourth row straight and shorter, terminating anteriorly at the level of buccal cirrus, DM_1 & 2_ short, three caudal cirri at the end of DK_1,2&4_; the generation time under laboratory conditions 12 ± 0.5 h, encystment frequent, conjugation observed.

Genus S*terkiella* Foissner *et al.* 1991

#### 16. *Sterkiella tetracirrata* Kumar *et al.,* 2015

Size in vivo 75–85 × 30–40 μm, after protargol impregnation about 75 × 30 μm, length to width ratio is 2:1; body oblong, firm to slightly flexible, pellicle, cytoplasm colourless; AZM extends 46% of the cell length comprising 35 membranelles; UM in *Stylonychia* pattern; four macronuclei, two to four micronuclei; 17 FVT cirri which includes three frontal, one buccal, four frontoventral, three post oral ventral, two pretransverse, four transverse cirri; one left and one right row of marginal cirri with an average of 25 and 28 cirri, respectively; six rows of dorsal kineties (DK_1–4_ and DM_1–2_); three caudal cirri, one each at the posterior ends of DK_1, 2 & 4_, placed equidistant; encystment observed frequently, conjugation not observed.

#### 17. *Sterkiella* sp

Size in vivo 80–90 × 35–45 μm, after protargol impregnation 80 × 36 μm, length to width ratio is 2.2:1; body firm to slightly flexible, pellicle, cytoplasm colourless; AZM extends 45% of the cell length comprising 28 membranelles; UM in *Stylonychia* pattern; two macronuclei, two to four micronuclei; 17 FVT cirri with three frontal, one buccal, four to five frontoventral, three to four postoral ventral, two pretransverse, three or four transverse cirri; one left and one right row of marginal cirri with about 16 and 17 cirri, respectively; six rows of dorsal kineties (DK_1-4_ and DM_1-2_); three caudal cirri, one each at the posterior ends of DK_1, 2 & 4_, placed equidistant; encystment observed frequently, conjugation not observed.

Genus *Tetmemena* Eigner, 1999

#### 18. *Tetmemena pustulata* (Müller, 1786) Ehrenberg, 1835

Size in vivo 100–120 × 40–50 μm, after protargol impregnation about 100 × 42 μm, length to width ratio around 2:1, body rigid with no cortical granules, body rigid with both the end rounded; AZM covers less than 50% of body length with an average of 36 adoral membranelles; UMs in *Stylonychia* pattern; two macronuclei, two micronuclei; 18 FVT cirri which include three frontal, one buccal, four frontoventral (arranged in oblique hook-shaped row), three postoral ventral placed equidistant to each other, two pretransverse, five transverse; one left and one right row of marginal cirri with 20 and 26 cirri, respectively; six rows of dorsal kineties (DK_1–4_ and DM_1, 2_), DK_4_ extends full body length, three caudal cirri not equidistant; generation time under laboratory conditions 8 h, encystment and conjugation infrequent.

#### 19. *Tetmemena saprai* Gupta *et al.,* 2020

Size in vivo 125–140 × 50–60 μm, after protargol impregnation about 100 × 45 μm, length to width ratio around 2:1, body rigid with no cortical granules, body lanceolate anteriorly and rounded posteriorly; AZM covers less than 50% of body length with an average of 42 adoral membranelles; UMs in *Stylonychia* pattern; two macronuclei, two to four micronuclei; 18 FVT cirri which include three frontal, one buccal, four frontoventral, three postoral ventral, two pretransverse, five transverse; one left and one right row of marginal cirri with 22 and 28 cirri, respectively; six rows of dorsal kineties (DK_1–4_ and DM_1, 2_) with DK_4_ shortened anteriorly, three caudal cirri not equidistant; generation time under laboratory condtions 8 h, encystment and conjugation infrequent.

#### 20. *Tememena vorax* Stokes, 1886

Size in vivo 90–100 × 40–50, after protargol impregnation about 87 × 40 μm, length to width ratio about 2:1; body ovoid, rigid, dorsoventrally flattened with a lanceolate anterior end and a tapering posterior end; AZM less than 50% of the cell length with an average of 34 membranelles; UM in *Stylonychia* pattern; two macronuclei, two micronuclei; 18 FVT cirri which include three frontal, one buccal, four frontoventral, three postoral ventral, two pretransverse, five transverse cirri characteristically arranged in two groups, a group of three cirri forming an oblique row (T_1_–T_3_), a detached group of two cirri reaching very close to the posterior border; one left and one right row of marginal cirri with 16 and 18 cirri, respectively; six rows of dorsal kineties (DK_1–4_ and DM_1, 2_), DK_1_-DK_3_ extend along the entire body length, the fourth kinety (DK_4_) somewhat shorter and the two dorsomarginals (DM_1 & 2_) terminate in the anterior half of the body, three caudal cirri barely distinguishable from the marginal; generation time under laboratory conditions 8 ± 0.5 h, encystment rare, conjugation very frequent.

#### 21. *Tetmemena* sp

Size in vivo 90–100 × 40–50 μm, after protargol impregnation about 91 × 43 μm, length to width ratio being 2:1; body rigid, dorsoventrally flattened, broad anterior end and a rounded tapering posterior end; AZM is less than 50% of the cell length with an average of 44 membranelles; UMs in *Stylonychia* pattern; two macronuclei, two micronuclei; 18 FVT cirri which include three frontal, one buccal, four frontoventral, posterior frontoventral cirri forms a small arc (III/2, IV/3 and VI/3), three postoral ventral, two pretransverse, five transverse cirri arranged in two groups of four and one, II/1,III/1, IV/1 and V1/1 arranged in an arc, V/1 separated towards the posterior end; one left and one right row of marginal cirri with 17 and 25 cirri, respectively; six rows of dorsal kineties (DK_1–4_ and DM_1 & 2_), DK_1_-DK_4_ extend along the entire body length, DM_1 & 2_ shortened posteriorly, three caudal cirri equidistant to each other; generation time under laboratory conditions 7 h, encystment frequent, conjugation rare.

Genus *Urosomoida* Hemberger in Foissner, 1982

#### 22. *Urosomoida* sp

Size in vivo 60–65 × 17–25 μm, after protargol impregnation about 58 × 17 μm; length to width ratio 3.6:1; body flexible, dorsoventrally flattened, cell broadest at the mid body, tapering posteriorly; AZM occupies about 30% of the body length with 16 membranelles; UMs in typical *Oxytricha* pattern; two macronuclei, two micronuclei; 16 FVT cirri with three frontal, one buccal, four frontoventral, three post oral ventral, two pretransverse, three transverse cirri, distance between II/1 and III/1 is more than the distance between III/1 and IV/1, closely placed tranverse cirri; one left and one right row of marginal cirri with an average of 13 and 16 cirri, respectively, both rows are wide open posteriorly; four rows of dorsal kineties (DK_1-3_ and DM_1_), DM_1_ shorter terminating in the anterior quarter of the cell; three caudal cirri, one each at the posterior ends of DK_1-3_, encystment and conjugation not observed.

Family Gonostomatidae Small & Lynn, 1985
Genus *Gonostomum* Sterki, 1878

#### 23. *Gonostomum* sp

Size in vivo 60–65 × 20–30 μm, after protargol impregnation about 62 × 23 μm on an average, length to width ratio 2.8:1; body flexible, ellipsoid, both ends more or less narrowly rounded; Oral apparatus in *Gonostonum* pattern, AZM covers about 42% of the body length with an average of 27 membranelles, UMs in *Gonostomum* pattern; two macronuclei, four micronuclei; three enlarged frontal cirri, one buccal cirrus, one short frontoterminal row with three to eight cirri, three rows of frontoventral cirri with first row having two to four cirri, second row having two to five cirri and third row having 6–10 cirri, two to six transverse cirri, one left and one right row of marginal cirri with 16 and 23 cirri on an average, respectively; three rows of dorsal kineties, three caudal cirri, one each at the posterior ends of DK_1–3_, encystment frequent, conjugation not observed.

Family Amphisiellidae Jankowski, 1979
Genus *Hemiamphisiella* Foissner, 1988

#### 24. *Hemiamphisiella terricola* Foissner, 1988

Size in vivo about 130–140 × 35–45 μm, after protargol impregnation is about 131 × 35 μm, length to width ratio 3.7:1; body flexible, outline sigmoidal, anterior portion slightly, posterior distinctly narrowed; AZM covers about 27% of body length with about 35 membranelles; UMs in optically intersecting; 19–27 macronuclei, one to six micronuclei; four frontal, one buccal, one amphisiellid median row with 30–46 cirri, one postoral ventral cirrus, one right ventral row having three to five cirri; one left and one right row of marginal cirri with 40 and 46 cirri on an average, respectively; three rows of dorsal kineties, three to four caudal cirri, one each at the posterior ends of each DK; encystment rare, conjugation not observed.

Family Holostichidae Fauré-Fremiet, 1961
Genus *Anteholosticha* Berger, 2003

#### 25. *Anteholosticha monilata* Kahl, 1932

Size in vivo about 146–176 × 35–39 μm, after protargol impregnation about 125 × 30 μm, length to width ratio is 4:1; body flexible, slender to ellipsoidal in shape, anterior end more or less narrowed, posterior broadly rounded; AZM covers about 28% of the body length with 41 membranelles; UMs optically intersecting; 13–18 macronuclei, three to seven micronuclei; three to six frontal cirri, one buccal, two or four frontoterminal cirri, 18–27 cirral pairs in mid ventral row of cirri, six to eight transverse cirri; one left and one right row of marginal cirri with an average of 41 cirri in each row; six rows of dorsal kineties, caudal cirri absent; encystment frequent, conjugation not observed.

Family Uroleptidae Foissner & Stoeck, 2008
Genus *Uroleptus* Ehrenberg, 1831

#### 26. *Uroleptus gallina* Müller, 1786

Size in vivo 125–145 × 40–52 μm, after protargol impregnation about 112 × 43 μm, length to width ratio is 2.6:1; body moderately flexible, slender, anterior end broad, posterior end narrowed giving a fish like appearance; AZM covers about 35% of the body length with an average of 44 membranelles; UMs intersecting and distinctly curved; two macronuclei, two micronuclei; one frontal cirrus, one buccal cirrus, one parabuccal cirrus, two to three frontoterminal cirri, midventral complex typically uroleptid composed of 23 pairs of cirri on an average with right row slightly larger than the left; long transverse cirral row composed of 12– 14 cirri; one left and one right row of marginal cirri with an average of 24 and 28 cirri, respectively; five rows of dorsal kineties, three caudal cirri, one each at the posterior ends of kineties DK_1, 2 & 3_; encystment frequent, conjugation not observed.

#### 27. *Uroleptus longicaudatus* Stokes, 1886

Size in vivo 160–170 × 40–50 μm, after protargol impregnation is 152 × 34 μm on an average, length to width ratio about 5:1; body fusiform, elongated, elliptical with a tail strongly retracted to about 12% of body length, body dorsoventrally flattened, cortical granules lacking, cytoplasm colorless; AZM occupies about 27% of body length with an average of 44 membranelles; UMs intersecting and distinctly curved; two macronuclei, two to four micronuclei; three frontal cirri, one buccal cirrus, two to three frontoterminal cirri, midventral complex typically uroleptid composed of 23 pairs of cirri, three to four transverse cirri; one left and one right row of marginal cirri with an average of 32 cirri on each row; five rows of dorsal kineties, three caudal cirri, one each at the posterior ends of kineties DK_1, 2 & 3_; encystment and conjugation rarely observed.

Order Urostylida Jankowski, 1979
Family Pseudokeronopsidae Borror & Wicklow, 1983
Genus *Pseudokeronopsis* Borror & Wicklow, 1983

#### 28. *Pseudokeronopsis* sp

Size in vivo 160–220 × 54–62 μm, after protargol impregnation about 165 × 44 μm, length to width ratio about 4:1; body highly flexible, elongated, shape often becomes plumper in culture, cell is brightly coloured, dark orange-browish pigment granules about 1 μm in size and invariably arranged in a ‘*rubra*-pattern’ (Song *et al.* 2006); AZM covers about 30% of the body length with an average of 50 membranelles; UMs in *pseudokeronopsid-*pattern; 52–78 macronuclei, two or three micronuclei; 12 pairs of frontal cirri, one buccal cirrus, two frontoterminal cirri, midventral complex with 28 pairs of cirri on an average with about two or three unpaired cirri at the posterior end, two pretransverse cirri, two to six transverse cirri; one left and one right row of marginal cirri with an average of 50 and 58 cirri, respectively; three rows of dorsal kineties (DK_1–3_), caudal cirri absent; encystment frequent, conjugation not observed.

Family Pseudourostylidae Jankowski, 1979
Genus *Pseudourostyla* Borror, 1972

#### 29. *Pseudourostyla cristata* Jerka-Dziadosz, 1964

Size in vivo about 190–250 × 60–90 μm, size after protargol impregnation 194 × 54 μm on an average, length to width ratio about 3.6:1; body flexible, slender, brownish, ellipsoidal with both ends broadly rounded, extrusomes present; AZM covers 37% of the body size with an average of 68 membranelles; UMs optically intersecting; 15–83 macronuclear nodules and two to nine micronuclei; one buccal cirrus, 18–22 frontal cirri, two or three frontoterminal cirri, midventral complex having 20–27 pairs of cirri, seven to nine transverse cirri, five rows of left and four rows of right marginal cirri; 8–10 rows of dorsal kineties, caudal cirri absent; encystment is frequent, conjugation not observed.

Family Urostylidae Bütschli, 1889
Genus *Diaxonella* Jankowski, 1979

#### 30. *Diaxonella trimarginata* Jankowski, 1979

Size in vivo about 120–150 × 40–70 μm, size after protargol impregnation 115 × 50 μm on an average, length to width ratio about 2:1; body flexible, slender, elongate ellipsoidal body with both ends rounded, cell is brightly colored with rose colored granules; AZM covers 36% of the body size with an average of 33 membranelles on an average; UMs optically intersecting; 50–67 macronuclei, 7–10 micronuclei; four to six buccal cirri, three frontal cirri, four frontal row cirri, two frontoterminal cirri, midventral complex having 15–26 pairs of cirri, two to four pretransverse, six to seven transverse cirri arranged in J-shaped row; one row of right marginal cirri with 40 cirri and four rows of left marginal cirri with an average of 24, 22, 20 and 21 cirri on each row (LMR_1–4_); three rows of dorsal kineties, caudal cirri absent; encystment frequent, conjugation not observed.

Genus *Urostyla* Ehrenberg, 1830

#### 31. *Urostyla grandis* Ehrenberg, 1830

Size in vivo about 240–260 × 120–140 μm, size after protargol impregnation 170–220 × 80–110 μm, length to width ratio about 2:1; body flexible, elongate body with both ends rounded, yellowish-green cortical granules; AZM covers 46% of the body length with an average of 45 membranelles; undulating membrane optically intersecting; numerous macronuclei with two to four micronuclei; eight frontal cirri, seven buccal cirri, three rows of parabuccal cirri with two to four, two and one cirri in each row, respectively, midventral complex having 17–18 pairs of cirri, seven transverse cirri, five rows of right marginal cirri with an average of 28 cirri in the innermost row and 38 cirri on the outermost row, four rows of left marginal cirri with an average of 25 cirri in the innermost row and 36 cirri on the outermost row; three rows of dorsal kineties, caudal cirri absent; encystment frequent, conjugation not observed.

Order Euplotida Small & Lynn, 1985
Family Aspidiscidae Ehrenberg, 1830
Genus *Aspidisca* Ehrenberg, 1830

#### 32. *Aspidisca* sp

Size in vivo about 40–60 × 20–30 μm, after protargol impregnation about 19 × 16 μm, length to width ratio about 1:1; small sized, ovoid, inflexible body with four dorsal ridges; two to four membranelles in anterior portion of adoral zone and 5–11 in posterior portion; one macronucleus horseshoe-shaped and one or two micronuclei; seven frontal cirri, five transverse cirri; four rows of dorsal kineties; encystment and conjugation not observed.

Family Euplotidae Ehrenberg, 1838
Genus *Euplotes* Ehrenberg, 1830

#### 33. *Euplotes aediculatus* Pierson, 1943

Size in vivo about 107–119 × 72–82 μm, after protargol impregnation 97 × 66 μm, length to width ratio about 1.5:1; body rectangular, rigid, dorso-ventrally flattened with inconspicuous dorsal grooves, pellicle colourless with refractive granules; macronucleus C-shaped with an arched back, one spherical micronucleus; AZM occupies 69% of the body length with an average of 44 membranelles; nine frontoventral cirri, five transverse cirri, two left marginal cirri, two caudal cirri, eight dorsolateral kineties, 20–24 pairs of basal bodies in the mid dorsolateral kinety (DK_5_), left and right most DK located towards the ventral side and are obviously short; double-eurystomus type of dargyrome on the dorsal surface and irregular on ventral side; encystment and conjugation not observed.

#### 34. *Euplotes woodruffi* Gaw, 1839

Size in vivo 100–132 × 68–82 μm, after protargol impregnations 95 × 66 μm, length to width ratio about 1.5:1, body ovoid, rigid, dorso-ventrally flattened with inconspicuous dorsal grooves; pellicle colourless with refractive granules, AZM occupies 79% of the body length with an average of 46 membranelles, preoral pouch in shape of inverted teardrop, invaginating the dorsal wall of buccal cavity; T-shaped macronucleus, one micronucleus; nine frontoventral cirri, five transverse cirri, two left marginal cirri, two caudal cirri, nine or ten dorsolateral kineties, 20–21 pairs of basal bodies in the mid dorsolateral kinety (DK_5_); double eurystomus type of dargyrome on the dorsal surface and irregular on ventral surface; encystment and conjugation not observed.

#### 35. *Euplotes* sp

Size in vivo about 49–52 × 40–46 μm, after protargol impregnation 30–45 × 19–36 μm, length to width ratio approximately 1.4:1; body oval, pellicle colourless; macronucleus C-shaped, one spherical micronucleus; adoral zone extends to 67% of body length, composed of 20–25 membranelles; 10 frontoventral, five transverse, two left marginal, two caudal cirri; seven dorsolateral kineties within six prominent dorsal ridges, mid dorsal kinety (DK_4_) with 11–16 dikinetids; dorsal silverline system of double eurystomus type and irregular on ventral; encystment frequent and conjugation not observed.

Class Heterotrichea Stein, 1859
Order Heterotrichida Stein, 1859
Family Blepharismidae Jankowski in Small & Lynn, 1985
Genus *Blepharisma* Perty, 1852

#### 36. *Blepharisma sinosum* Sawaya, 1940

Size in vivo 100–120 × 30–40 μm, size after protargol impregnation about 80 × 30 μm, length to width ratio about 4.7:1, body spindle-shaped with tapered anterior end, pellicle flexible with numerous pink cortical granules arranged in five to seven longitudinal rows between the kineties; S-shaped AZM occupies 50% of the body length with an average of 45 membranelles; moniliform macronucleus with four to seven nodules connected by nuclear bridges, macronuclear nodule shape varies from spherical to ellipsoidal with 10–20 spherical micronuclei dispersed throughout the cytoplasm; 20–30 somatic kineties; encystment and conjugation observed.

#### 37. *Blepharisma undulans* Stein, 1867

Size in vivo 100–120 × 30–40 μm, after protargol impregnation about 90 × 30 μm, length to width ratio 3:1; body flexible, spindle shape, elongate, lenticular with anterior conspicuously pointed, posterior end widely rounded, contains numerous pink cortical granules distributed in numerous longitudinal rows between kineties, peristome narrow extending to mid-body region, AZM occupies 50% of the body length with an average of 58 membranelles, paroral membrane composed of dikinetids; two ellipsoidal macronuclear nodules, one located in anterior and the other in posterior half of body connected by a very thin strand, the micronuclei are few in number varying from two to six; 20–30 somatic kineties arranged longitudinally; encystment and conjugation observed.

#### 38. *Blepharisma* sp

Size in vivo 80–90 × 20–30 μm, after protargol impregnation about 60 × 20 μm, length to width ratio 3:1; body slender, flexible and slightly bilaterally flattened with slightly pale pink pigmentation, anterior end bluntly pointed, posterior broadly rounded; peristome extends almost half the body length; contractile vacuole located posteriorly, AZM occupies 65% of the body length with an average of 36 membranelles; paroral membrane inconspicuous; single spherical to ovoid macronucleus located at the mid-body region, two or three spherical micronuclei associated with the macronucleus; contains 10–20 somatic longitudinal kineties; encystment and conjugation observed.

Family Spirostomidae Stein, 1867
Genus *Spirostomum* Ehrenberg, 1833

#### 39. *Spirostomum minus* Roux, 1901

Size in vivo of fully extended organism 400–800 × 80–85 μm, size in vivo of contracted organism 175–262 × 55–115 μm, length to width ratio about 3:1 (contracted organism) to 7:1 (fully extended organism); body flexible, elongated, vermiform, slender, anterior and posterior end tapering; yellow-brown in color when observed under low magnification due to the presence of pale brown cortical granules, contractile vacuole at the posterior end along the dorsal side; moniliform macronucleus as a chain of beads, consists of usually 9–25 ellipsoidal beads with 12 micronuclei; AZM reaches to the body center about 50% of the body with a length of about 112 μm length; somatic ciliature composed of about 30–40 spiral ciliary kineties; encystment and conjugation not observed.

Family Stentoridae Carus, 1863
Genus *Stentor* Oken, 1815

#### 40. *Stentor* sp

Size in vivo 230–300 × 130–140 μm, after protargol impregnations 190 × 105 μm, length to width ratio about 1.7:1; body large, trumpet or horseshoe shaped, broad anteriorly and tapering at the posterior end, cortical granules intensively blue green, ring of cilia spiraling around the anterior end; single moniliform macronucleus with several micronuclei; AZM present at the margin in shape of elliptical to circular forming a small buccal cavity which leads to the cytostome with 150 membranelles on average; encystment observed, conjugation not observed.

Order Loxodida Jankowski, 1980
Family Loxodidae Bütschli, 1889
Genus *Loxodes* Ehrenberg, 1830

#### 41. *Loxodes* sp

Size in vivo 144–155 × 70–80 μm, after protargol impregnations 130 × 68 μm, length to width ratio 2:1; body elongated with pointed ends, anterior portion forming beak like structure, lanceolate body shape; colourless to light brownish cortical granules present, contractile vacuoles absent; buccal field occupying 1/6–1/7^th^ of body length, have an elongated bursiform outline; four or five macronuclei and micronuclei; somatic cilia are approximately 5 μm in length, right somatic ciliary rows number about 11–13; encystment not observed, conjugation observed.

Class Litostomatea Small & Lynn, 1981
Order Dileptida Jankowski, 1978
Family Dileptidae Jankowski, 1980
Genus *Dileptus* Dujardin, 1841

#### 42. *Dileptus* sp

Size in vivo 147–155 × 26–30 μm, after protargol impregnations 136 × 25 μm, length to width ratio 5.7:1, rigid body divided into two main parts: anteriorly located slender elongation called proboscis and trunk, body distinctly cylindrical, widest mid-body tapering slightly at the tail and proboscis, dark cytoplasm with several contractile vacuoles, large number of spherical to ovoid macronuclei containing numerous small and spherical micronuclei, body except for the oral region is covered with longitudinal rows of somatic cilia; encystment observed, conjugation not observed.

Order Haptorida Corliss, 1974
Family Lacrymariidae de Fromentel, 1876
Genus *Lacrymaria* Bory, 1826

#### 43. *Lacrymaria* sp

Size in vivo 120–150 × 20–30 μm, size after protargol impregnations 100 × 10 μm, length to width ratio 6:1, tear drop shaped body with small head and a long, active, slender, and flexible neck, the body is covered with cilia except the apical zone which remains unciliated, posterior end is bluntly pointed; encystment and conjugation not observed.

Order Pleurostomatida Schewiakoff, 1896
Family Litonotidae Kent, 1882
Genus *Loxophyllum* Dujardin, 1840

#### 44. *Loxophyllum* sp

Size in vivo 135–145 × 45–55 μm, after protargol impregnations 125 × 45 μm, length to width ratio 3:1, broad leaf-shaped body with a beak-like anterior end and a blunt posterior end, body laterally compressed, contractile vacuoles located in the posterior half of the body, numerous colourless cortical granules; two ellipsoidal macronuclear nodules; four to six conspicuous longitudinal ridges; encystment and conjugation not observed.

Genus *Litonotus* Wrzesniowski, 1870

#### 45. *Litonotus* sp

Size in vivo 205–215 × 60–70 μm, after protargol impregnations 160 × 50 μm, length to width ratio 3:1; body laterally flattened, flexible, lanceolate, pointed posterior end, spindle-shaped with long neck bent towards the dorsal side, colourless cortical granules; four ellipsoidal macronuclear nodules; mouth located along the anterior margin extending from anterior pointed pole to near the middle of the cell; encystment and conjugation not observed.

Order Spathidiida Foissner & Foissner, 1988
Family Spathidiidae Kahl in Doflein & Reichenow, 1929
Genus *Spathidium* Dujardin, 1841

#### 46. *Spathidium* sp

Size in vivo 170–180 × 30–40 μm, after protargol impregnations 150 × 25 μm, length to width ratio is about 6:1; body flexible, elongated, posteriorly blunt or rounded, laterally compressed; prominent anterior oral bulge, contractile vacuole single and posteriorly located; single spherical macronucleus with several micronuclei; uniformly ciliated with about 20 somatic kineties; encystment and conjugation not observed.

Class Phyllopharyngea de Puytorac *et al.,* 1974
Order Chlamydodontida Deroux, 1976
Family Chlamydodontidae Stein, 1859
Genus *Chilodonella* Strand, 1928

#### 47. *Chilodonella* sp

Size in vivo 45–55 × 25–35 μm, after protargol impregnations 30 × 23 μm, length to width ratio 1.7:1; body dorsoventrally flattened, oval in shape, distinct pre-oral beak at the anterior end cytoplasm transparent; the cell has a flat ventral surface and an arched dorsal surface, the ventral surface is ciliated with many number of somatic kineties placed longitudinally, the dorsal surface is unciliated; single spherical to ellipsoidal macronucleus located near the posterior end with two or three micronuclei; encystment and conjugation not observed.

Class Oligohymenophorea de Puytorac *et al.,* 1974
Order Peniculida Fauré-Fremiet in Corliss, 1956
Family Parameciidae Dujardin, 1840
Genus *Paramecium* O. F. Müller, 1773

#### 48. *Paramecium multimicronucleatum* Powers & Mitchell, 1910

Size in vivo 204–210 × 55–65 μm, after protargol staining 196 × 53 μm, length to width ratio 4:1, body slipper-shaped with cilia covering the entire body, trichocyst present; AZM occupies 40% of the body with an average of 23 membranelles; encystment not observed, conjugation observed.

Family Frontoniidae Kahl, 1926
Genus *Frontonia* Ehrenberg, 1833

#### 49. *Frontonia* sp

Size in vivo 300–350 × 200–250 μm, after protargol impregnations 220 × 130 μm on an average, length to width ratio 1.5:1; ovoid with an oral depression in the anterior half of the cell, flexible, dorsoventrally flattened with anterior end broad and posterior end narrowed; cytoplasm colourless; single ellipsoidal macronucleus centrally located; uniformly ciliated with several somatic kineties; encystment and conjugation not observed.

Order Pleuronematida Fauré-Fremiet in Corliss, 1956
Family Cyclidiidae Ehrenberg, 1838
Genus *Cyclidium* O. F. Müller, 1773

#### 50. *Cyclidium* sp

Size in vivo 20–30 × 13–15 μm, after protargol staining is about 14 × 9 μm, body length to width ratio 1.5:1; body elongated-ovoid, ventral side almost straight with dorsal evenly convex, apical end free from cilia, anterior end large and flat, posterior end broadly rounded; cytoplasm colorless; prominent undulating membrane on its right margin, paroral membrane gently curved extending posteriorly to about 3/4^th^ of cell length; two macronuclear nodules, spherical in shape and adjacent to each other, one micronucleus; 11–12 somatic kineties arranged longitudinally; contains single caudal cilium of length 20 μm; encystment and conjugation not observed.

Order Loxocephalida Jankowski, 1964
Family Loxocephalidae Jankowski, 1964
Genus *Dexiotricha* Stokes, 1885

#### 51. *Dexiotricha* sp

Size in vivo 55–65 × 25–35 μm, after protargol staining is about 45 × 20 μm; body length to width ratio 2:1; body shape long ellipsoidal, with ventral side straight, dorsal side gently curved elongated ovoid ciliates; cytoplasm colourless and containing numerous granules of about equal size; single macronucleus present near mid-body; contains single flexible and extremely long caudal cilium; encystment observed and conjugation not observed.

Order Sessilida Kahl, 1933
Family Vorticellidae Ehrenberg, 1838
Genus *Vorticella* Linnaeus, 1767

#### 52. *Vorticella* sp

Size in vivo 55–80 × 40–50 μm, after protargol impregnations 45 × 30 μm, length to width ratio 1.3:1; inverted bell shaped body with a long coiled stalk of length 100 μm and diameter of 3–4 μm; central part of cell is filled with refractile reserve granules; single long and worm-like macronucleus with a single spherical micronucleus; contains ring of cilia on the oral end; encystment and conjugation not observed.

Class Prostomatea Schewiakoff, 1896
Order Prorodontida Corliss, 1974
Family Colepidae Ehrenberg, 1838
Genus *Coleps* Nitzsch, 1817

#### 53. *Coleps* sp

Size in vivo 110–120 × 60–70 μm, after protargol impregnations 90 × 40 μm body length to width as 1.5:1, barrel shaped body, typically covered with spikes, consists of calcium carbonate cage, anterior end broad and posterior end moderately rounded; AZM short and obliquely arranged; one macronucleus, one micronucleus; longitudinally placed 20–25 somatic kineties; encystment not observed, conjugation frequent.

Class Colpodea Small & Lynn, 1981
Order Colpodida de Puytorac *et al.,* 1974
Family Colpodidae Bory de St. Vincent 1826
Genus *Colpoda* O. F. Müller, 1773

#### 54. *Colpoda magna* Gruber, 1880

Size in vivo 110–120 × 125 μm, after protargol impregnations 100 × 70 μm, body length to width 2:1; body having shape of bean or kidney, the anterior body broad and blunt and the posterior half tapering; colorless protoplasm with mass of black granules at the posterior end, endoplasm contains multiple food vacuoles; one macronucleus, one micronuclei; encystment and excystment frequent, conjugation rare.

#### 55. *Colpoda* sp

Size in vivo 40–50 × 30–40 μm, size after protargol impregnations 30 × 20 μm, body length to width ratio 1.3:1; body ellipsoidal to broadly ellipsoidal with distinct concavity at the oral opening, outline usually distinctly bulged by a mass of food vacuoles; one single ellipsoidal macronucleus with a single micronucleus attached; 15 ciliary rows arranged longitudinally; encystment and excystment frequent, conjugation rare.

## Conclusion

This documentation of freshwater ciliates is first of its kind from Delhi, India. Majority of the ciliates reported in this study belong to class Spirotrichea exhibiting maximum ciliate diversity with 23 genera and 35 species. Ciliates belonging to other classes have less representation i.e. class Heterotrichea with four genus and six species, class Litostomatea and Oligohymenophorea with five genus and five species, class Phyllopharyngea and Prostomatea with only one genus and one species, and class Colpodea with one genus and two species. Maximum ciliate diversity was obtained from Okhla bird sanctuary site (40 species) and least was from Rithala site (1 species). Most common species were *Aponotohymena australis, Oxytricha granulifera, Tetmemena saprai, Paraurostyla coronata, Urosomoida* sp. and *Gastrostyla* sp. present in more than five sites. Rest of the species were rarely observed and present in less than five sites.

Eight ciliate species belonging to the class Spirotrichea mentioned in this study, namely, *Aponotohymena isoaustralis* (Gupta *et al.* 2017)*, Architricha indica* (Gupta *et al.* 2006)*, Paraurostyla coronata* (Arora *et al.* 1999)*, Rubrioxitricha indica* (Naqvi *et al.* 2006)*, Stylonychia ammermanni* (Gupta *et al.* 2001), and *Tetmemena saprai* (Gupta *et al.* 2020) were reported for the first time from Delhi region. This study indicates that the freshwater bodies in Delhi exhibits rich ciliate diversity. This work has given us a lead to further explore and document the diverse ciliate fauna from different parts of India.

## Acknowledgements

The authors appreciate the facilities provided by the Principal, Acharya Narendra Dev College, University of Delhi for carrying out the present study. The support extended by the Principal, Maitreyi College, University of Delhi is thankfully acknowledged. We thank the Department of Zoology, University of Delhi for providing instrumental facilities. We also thank AIRF, JNU for assistance in DIC microscopy. The work was supported by Senior Research Fellowships to JSA and SM from CSIR and to SS from UGC, New Delhi, India.

## References

Abraham, J.S., Sripoorna, S., Choudhary, A., Toteja, R., Gupta, R., Makhija, S. & Warren, A. (2017) Assessment of Heavy Metal Toxicity in Four Species of Freshwater Ciliates (Spirotrichea; Ciliophora) from Delhi, India. Current Science 113, 2141–2150.

Abraham, J.S., Sripoorna, S., Dagar, J., Jangra, S., Kumar, A., Yadav, K., Singh, S., Goyal, A., Maurya, S., Gambhir, G., Toteja, R., Gupta, R., Singh, D.K., El-Serehy, H.A., Al-Misned, F.A., Al-Farraj, S.A., Al-Rasheid, K.A., Maodaa, S.A. & Makhija, S. (2019a) Soil ciliates of the Indian Delhi Region: their community characteristics with emphasis on their ecological implications as sensitive bio-indicators for soil quality. Saudi Journal of Biological Sciences, 26, 1305–1313.

Abraham, J.S., Sripoorna, S., Maurya, S., Makhija, S., Gupta, R. & Toteja, R. (2019b) Techniques and tools for species identification in ciliates: a review. International Journal of Systematics and Evolutionary Microbiology, 69, 877–894.

Adl, S.M., Bass, D., Lane, C.E., Lukeš, J., Schoch, C. L., Smirnov, A., Agatha, S., Berney, C., Brown, M.W., Burki, F., Cardenas, P., Cepicka, I., Chistyakova, L., Campo, J.D., Dunthorn, M., Edvardsen, B., Eglit, Y., Guillou, L., Hampl, V., Heiss, A.A., Hoppenrath, M., James, T.Y., Karnkowska, A., Karpov, S., Kim, E., Kolisko, M., Kudryavtsev, A., Lahr, D.J.G, Lara, E., Gall, E., Lynn, D.H., Mann, D.G., Massana, R., Mitchell, E.A.D., Morrow, C., Park, J.S., Pawlowski, J.W., Powell, M.J., Ritcher, D.J., Rueckert, S., Shadwick, L., Shimano, S., Spiegel, F.W., Torruella, G., Youssef, N., Zlatogursky, V., Zhang, Q. (2019) Revisions to the Classification, Nomenclature, and Diversity of Eukaryotes. Journal of Eukaryotic Microbiology, 65, 623–649.

Aghaindum, G.A. & Menbohan, S.F. (2012) Determination of optimum period for ciliated protozoa colonizing of an artificial substrate in a tropical aquatic ecosystem. International Journal of Environmental Science and Technology, 9, 655–660.

Ammermann, D., & Schlegel, M. (1983). Characterization of two sibling species of the Genus *Stylonychia* (Ciliata, Hypotricha): *S. mytilus* Ehrenberg, 1838 and *S. lemnae* n. sp. I. Morphology and reproductive Behavior. The Journal of Protozoology, 30, 290–294.

Arora, S., Gupta, R., Kamra, K. & Sapra, G.R. (1999) Characterization of *Paraurostyla coronata* sp. n. including a comparative account of other members of the genus. Acta Protozoologica, 38, 133–144.

Basuri, C.K., Pazhaniyappan, E., Munnooru, K., Chandrasekaran, M., Vinjamuri, R.R., Karri, R. & Mallavarapu, R.V. (2020) Composition and distribution of planktonic ciliates with indications to water quality in a shallow hypersaline lagoon (Pulicat Lake, India). Environmental Science and Pollution Research, 27, 18303–18316.

Berger, H. (1999) Monograph of Oxytrichidae (Ciliophora, Hypotrichia). Monographiae Biologicae. Springer, 78, 1–1080.

Bharti, D. & Kumar, S. (2019) Ten new records of Protozoan ciliates (Protozoa: Ciliophora) from India. Records of the Zoological Survey of India, 119, 111–119.

Bharti, D., Kumar, S., La Terza, A. & Chandra, K. (2019) Morphology and ontogeny of *Tetmemena pustulata indica* nov. subspec. (Ciliophora, Hypotricha), from the Thane Creek, Mumbai, India. European Journal of Protistology, 71, 125629.

Bhatia, B. L. (1936) The Fauna of British India including Ceylon and Burma. Protozoa: Ciliophora. Taylor and Francis, London, 498 pp.

Bindu, L., Purushothaman, J., Das, A.K., Nandi, N.C. & Kumar, S. (2018) Protozoa (Free-living). *In*: Chandra, K., Gupta, D., Gopi, K.C., Tripathy, B. & Kumar, V., Faunal diversity of Indian Himalaya. Zoological Survey of India, Kolkata, pp. 45–57.

Chanda, S., Barman, G.D. & Bandyopadhyay, P.K. (2019) A checklist of trichodinid ciliates (Ciliophora: Peritrichida: Trichodinidae) from India. Records of the Zoological Survey of India, 119, 424–437.

Chapman-Andresen, C. (1958) Pinocytosis of inorganic salts by *Amoeba proteus* (C*haos diffluens*). Comptes rendus des travaux du Laboratoire Carlsberg, Série chimique, 31, 77–92.

Corliss, J.O. (1979) The Ciliated Protozoa: Characterization, Classification and Guide to the Literature. Vol. 2. Pergamon Press, Oxford, 472 pp.

Corliss, J.O. (2002) Biodiversity and biocomplexicity of the Protists and an overview of their significant roles in maintenance of our biosphere. Acta Protozoologica, 41, 199–219.

Debastiani, C., Meira, B.R., Lansac-Tȏha, F.M., Velho, L.F.M., Lansac-Tȏha, F.A. (2016) Protozoa ciliates community structure in urban streams and their environmental use as indicators. Brazilian Journal of Biology, 76, 1043–1053.

Dopheide, A., Lear, G., Stott, R. & Lewis, G. (2008) Molecular characterization of ciliate diversity in stream biofilms. Applied and Environmental Microbiology, 74, 1740–1747.

Ehrenberg, C.G. (1835) Das Leuchten des Meeres: neue Beobachtungen nebst Übersicht der Hauptmomente der geschichtlichen Entwicklung dieses merkwürdigen Phänomens. Gedruckt in der Druckerei der Königlichen Akademie der Wissenschaften. Berlin, 176 pp. (in German).

Ehrenberg, C.G. (1830) Beiträge zur Kenntniß der Organisation der Infusorien und ihrer geographischen Verbreitung, besonders in Sibirien. Abhandlungen der Preussischen Akademie der Wissenschaften, 1830, 1–88.

Elangovan, S.S. & Gauns, M. (2018) A checklist of tintinnids (loricate ciliates) from the coastal zone. Environmental Monitoring and Assessment, 190, 1–19.

Engelmann, T.W. (1862) Zur Naturgeschichte der Infusionsthiere. Zeitschrift fűr wissenschaftliche Zoologie, 11, 347–393.

Fenchel T. (1987) Ecology of Protozoa: The Biology of Free-Living Phagotrophic Protists. Science Tech Publishers, Madison (WI), 197 pp.

Feulgen, R. & Rossenbeck, H. (1924) Mikroskopisch-chemischer Nachweis einer Nucleinsaure vom Typus der Thymonucleinsaure und die darauf beruhende elektive Farbung vom Zellkernen in mikroskopischen Praparaten. Zeitschrift fur Physiologische Chemie, 135, 203–248 (in German).

Finlay, B.J., Maberly, S.C. & Cooper, J.I. (1997) Microbial diversity and ecosystem function. OIKOS, 80, 209–213.

Foissner, W. (1998) An updated compilation of world soil ciliates (Protozoa, Ciliophora) with ecological notes, new records, and descriptions of new species. European Journal of Protistology, 34, 195–235.

Foissner, W. & O’donoghue, P.J. (1989) Morphology and infraciliature of some freshwater ciliates (Protozoa: Ciliophora) from Western and South Australia. Invertebrate Systematics, 3, 661–696.

Foissner, W. (1988) Gemeinsame Arten in der terricolen Ciliatenfauna (Protozoa: Ciliophora) von Australien und Afrika. Stapfia, Linz, 17, 85–133.

Foissner, W. & Adam, H. (1983) Morphologie und Morphogenese des Bodenciliaten Oxytricha granulifera sp. n. (CiIiophora, Oxytrichidae). Zoologica Scripta, 12, 1–11.

Gaw, Z.H. (1939) *Euplotes woodruffi* sp. nov. Archiv für Protistenkunde, 93, 1–5.

Gelei, J. & Szabados, M. (1950) Tömegprodukció városi esövizpocsolyában (Massenproduktion in einer städtischen Regenwasserpfütze). Annales biologicae Universitatum Szegediensis, 1, 249–294.

Groliére, C.A., Chakli, R., Sparagano, O. & Pepin, D. (1990) Application de la colonisation d’un substratartificiel par les Ciliés à l’Étude de la qualité des eauxd’unerivière. European Journal of Protistology, 25, 381–390.

Gruber, A. (1880) Neue Infusorien. Zeitschrift für wissenschaftliche Zoologie, 33, 439–466.

Gupta, R., Abraham, J.S., Somasundaram, S., Toteja, R., Makhija, S., El-Serehy, H.A. (2017). Taxonomic and morphogenetic description of the freshwater ciliate *Aponotohymena isoaustralis* n. sp. (Ciliophora; Oxytricha) isolated from Sanjay Lake, Delhi, India. Acta Protozoologica, 56, 93–107.

Gupta, R., Abraham, J.S., Sripoorna, S., Maurya, S., Toteja, R., Makhija, S., Al-Misned, F. A., El-Serehy, H.A. (2020) Description of a new species of *Tetmemena* (Ciliophora, Oxytrichidae) using classical and molecular markers. Journal of King Saud University-Science, 32, 2316–2328.

Gupta, R., Arora, S., Kamra, K. & Sapra, G.R. (2002) Biodiversity of Genus *Oxytricha* in the Delhi Region of the Indian Subcontinent. Japanese Journal of Protistology, 35, 42.

Gupta, R., Kamra, K. & Sapra, G.R. (2006) Morphology and cell division of the oxytrichids *Architricha indica* nov. gen., nov. sp. and *Histriculus histrio* (Müller, 1773), Corliss, 1960 (Ciliophora, Hypotrichida). European Journal of Protistology, 42, 29–48.

Gupta, R., Kamra, K., Arora, S. & Sapra, G.R. (2001) *Stylonychia ammermanni* sp. n. a new oxytrichid (Ciliophora, Hypotrichida) ciliate from the river Yamuna, Delhi, India. Acta Protozoologica, 40, 75–82.

Gupta, R., Kamra, K., Arora, S. & Sapra, G.R. (2003) *Pleurotricha curdsi* (Shi, Warren and Song 2002) nov. comb. (Ciliophora: Hypotrichida): morphology and ontogenesis of an Indian population; redefinition of the genus. European Journal of Protistology, 39, 275–285.

Gupta, R., Makhija, S., Toteja, R. (2018). Cell Biology Practical Manual. Prestige Publishers, Delhi.

Hou, C., Shi, X., Liu, G., Liu, S., Zhu, X. & Xu, H. (2016) Use of protozoa for assessing water quality in a humid subtropical Urban wetland ecosystem, southern china. Environment Pollution and Climate Change, 1, 105.

Jankowski, A.W. (1979) Revision of the order Hypotrichida Stein, 1859. Generic catalogue, phylogeny, taxonomy. Trudy Zoologitscheskogo Instituta, Leningrad, 86, 48–85.

Jerka-Dziadosz, M. (1964) *Urostyla cristata* sp. n. (Urostylidae, Hypotrichida): the morphology and morphogenesis. Acta Protozoologica, 2, 123–128.

Kahl, A. (1932) Urtiere oder Protozoa. I: Wimpertiere oder Ciliata (Infusoria), 3. Spirotricha. Tierwelt Deutschlands und der angrenzenden Meeresteile, 25, 399–650.

Kalavati, C. & Raman, A.V. (2008) Taxonomy and Ecology of Ciliated Protozoa from Marginal Marine Environments of East Coast of India. Records of the Zoological Survey of India, Occassional Paper no. 282, 1–136.

Kamra, K. & Kumar, S. (2010) *Notohymena saprai* sp. nov, a new oxytrichid ciliate (Protozoa, Ciliophora) from the Valley of Flowers, a Himalayan bioreserve region; description and morphogenesis of the new species. Indian Journal of Microbiology, 50, 33–45.

Kamra, K., Sapra, G.R. & Ammermann, D. (1994) *Coniculostomum bimarginata* n. sp., a new hypotrich ciliate: description and systematic relationships. European Journal of Protistolology, 30, 55–67.

Kaur, H., Shashi, Negi, R.K. & Kamra, K. (2019) Morphological and molecular characterization of *Neogastrostyla aqua* nov. gen., nov. spec. (Ciliophora, Hypotrichia) from River Yamuna, Delhi; comparison with *Gastrostyla-*like genera. European Journal of Protistology, 68, 68–79.

Kumar, S., Kamra, K., Bharti, D., La Terza, A., Sehgal, N., Warren, A. & Sapra, G.R. (2015) Morphology, morphogenesis, and molecular phylogeny of *Sterkiella tetracirrata* n. sp. (Ciliophora, Oxytrichidae), from the Silent Valley National Park, India. European Journal of Protistolology, 51, 86–97.

Li, J., Liao, Q., Li, M., Zhang, J., Tam, N.F. & Xu, R. (2010) Community Structure and Biodiversity of Soil Ciliates at Dongzhaigang Mangrove Forest in Hainan Island, China. Applied and Environmental Soil Science, 2010, 1–8.

Lynn, D.H. (2008) The ciliated protozoa: Characterization, classification and guide to the literature (3^rd^ ed.). Netherlands: Springer Science & Business Media.

Lynn, D.H. & Small, E.B. (2000) Phylum Ciliophora. In: J. J. Lee, G. F. Leedale & P. Bradbury (Eds.), An Illustrated Guide to the Protozoa, (2nd ed.), 371–656.

Madoni, P. (2000) The acute toxicity of nickel to freshwater ciliates. Environmental Pollution, 109, 53–59.

Mahajan, K.K. & Nair, K.N. (1971) On some freshwater ciliates (Protozoa) from Calcutta and its environs. Records of the Zoological Survey of India, 63, 1–4.

Makhija S., Abraham J.S., Somasundaram S., Toteja R. & Gupta R. (2016) Mapping the hidden diversity of free living fresh water ciliates from Delhi region, India. Protistology, 10, 43.

Mironova E., Telesh I., Skarlato S. (2012) Diversity and seasonality in structure of ciliate communities in the Neva Estuary (Baltic Sea). Journal of Plankton Research, 34, 208–220.

Müller, O.F. (1786) Animalcula Infusoria Fluviatilia et Marina, quae Detexit, Sytematice Descripsit et ad Vivum Delineari Curavit. N. Mölleri, Hauniae, 367p. (in Latin).

Naqvi, I., Gupta, R., Borgohain, P. & Sapra, G.R. (2006) Morphology and morphogenesis of *Rubrioxytricha indica* n. sp. (Ciliophora: hypotrichida). Acta Protozoologica, 45, 53–64.

Naqvi, I., Gupta, R., Makhija, S., Toteja, R., Abraham, J.S., Sripoorna, S., El-Serehy, H.A. & Al-Misned, F.A. (2016) Morphology and morphogenesis of a new oxytrichid ciliate, *Notohymena limus* n. sp. (Ciliophora, Oxytrichidae) from Delhi, India. Saudi Journal of Biological Sciences, 23, 789–794.

Pierson, B.F. (1943) A comparative morphological study of several species of *Euplotes* closely related to *Euplotes patella*. Journal of Morphology, 72, 125–165.

Powers, J.H. & Mitchell C. (1910) A new species of *Paramecium* (*P. multimicronucleata*) experimentally determined. Biological Bulletin, 19, 324–332.

Purushothaman, J., Bindu, L., Makhija, J., Toteja, R. & Gupta, R. (2017) Protozoa: Ciliophora (Ciliates). In K. Chandra, K.C. Gopi, D.V. Rao, K. Valarmathi and J.R.B. Alfred (Eds.), Current Status of Freshwater Faunal Diversity in India (pp 37–54). Zoological survey of India, Kolkata.

Rakshit, D., Sarkar, S.K. (2016) Diversity, distribution and polymorphism of loricate ciliate tintinnids along Hoogly estuary, India. Journal of the Marine Biological Association of India, 58, 61–68.

Roux, J. (1901) Faune infusorienne des eaux stagnantes des environs de Genève. Mémoires de l’Institut National Genevois, 19, 1–148.

Sawaya, M.P. (1940) Sobre um ciliado novo de São Paulo: *Blepharisma sinuosum* sp. n. (Ciliata, Heterotricha) e sobre a sub-ordem Odontostomata, nom. nov. Boletim da Faculdade de Filosofia Ciências e. Letras Universidade de São Paulo, 19, 303–308.

Shi, X., Warren, A., & Song, W. (2002) Studies on the morphology and morphogenesis of *Allotricha curdsi* sp. n. (Ciliophora: Hypotrichida). Acta Protozoologica, 41, 397–406.

Shukla, U. & Gupta, P.K. (2001) Assemblage of ciliated protozoan community in a polluted and non-polluted environment in a tropical lake of central Himalaya: Lake Naini Tal, India. Journal of Plankton Research, 23, 571–584.

Singh, J. & Kamra, K. (2013) *Paraurosomoida indiensis* gen. nov., sp. nov., an oxytrichid (Ciliophora, Hypotricha) from Kyongnosla Alpine Sanctuary, including note on non-oxytrichid Dorsomarginalia. European Journal of Protistology, 49, 600–610.

Singh, J. & Kamra, K. (2015) Molecular phylogeny of *Urosomoida agilis*, and new combinations: *Hemiurosomoida longa* gen. nov., comb. nov., and *Heterourosomoida lanceolata* gen. nov., comb. nov. (Ciliophora, Hypotricha). European Journal of Protistology, 51, 55–65.

Sivasankar, R., Ezhilarasan, P., Kumar, S., Naidu, S.A., Rao, G.D., Kanuri, V.V., Rao, V.R. & Ramu, K. (2018) Loricate ciliates as an indicator of eutrophication status in the estuarine and coastal waters. Marine Pollution Bulletin, 129, 207–211.

Somasundaram, S., Abraham, J.S., Gupta, R., Makhija, S. & Toteja, R. (2015) Diverse freshwater Spirotrich ciliate fauna from Okhla Bird Sanctuary, Delhi, India. Global Journal for Research Analysis, 4, 37–41.

Somasundaram, S., Abraham, J.S., Maurya, S., Toteja, R., Gupta, R. & Makhija, S. (2019) Expression and molecular characterization of stress-responsive genes (*hsp70* and *Mn-sod*) and evaluation of antioxidant enzymes (CAT and GPx) in heavy metal exposed freshwater ciliate, *Tetmemena* sp. Molecular Biology Reports, 46, 4921–4931.

Song, W., Warren, A., Roberts, D., Wilbert, N., Li, L., Sun, P., Hu, X. & Ma, H. (2007). Comparison and redefinition of four marine, coloured *Pseudokeronopsis* spp. (Ciliophora: Hypotrichida), with emphasis on their living morphology. Acta Protozoologica, 45, 271.

Stein, F. (1859) Der Organismus der Infusionstheire nach eigenen Forschungen insystematischer Reithenfolgebearbeitet I. Abtheilung. Algemeiner Theil und Naturges chichchte der hypotrichen Infusionsthiere. W. Engelmann, Leipzing, 206 pp.

Stein, F. (1867) Der Organismus der Infusionsthiere nach eigenen Forschungen in systematischer Reihenfolge bearbeitet. II Abtheilung. 1) Darstellung der neuesten Forschungsergebnisse über Bau, Fortpflanzung und Entwickelung der Infusionsthiere. 2) Naturgeschichte der heterotrichen Infusorien. Wilhelm Engelmann, Leipzig, 355 pp.

Stokes, A.C. (1886) Some new hypotrichous infusoria. Proceedings of the American Philosophical Society, 23, 21–30.

Tarbe, A.L., Unrein, F., Stenuite, S., Piriot, S., Sarmento, H., Sinyinza, D. & Descy, J.P. (2011) Protist herbivory: a key pathway in the pelagic foodweb of lake Tanganyika. Microbial Ecology, 62, 314–323.

Toteja, R., Makhija, S., Sripoorna, S., Abraham, J.S. & Gupta, R. (2017) Influence of copper and cadmium toxicity on antioxidant enzyme activity in freshwater ciliates. Indian Journal of Experimental Biology, 55, 694–701.

